# Interpretable machine learning framework reveals novel gut microbiome features in predicting type 2 diabetes

**DOI:** 10.1101/2020.04.05.024984

**Authors:** Wanglong Gou, Chu-wen Ling, Yan He, Zengliang Jiang, Yuanqing Fu, Fengzhe Xu, Zelei Miao, Ting-yu Sun, Jie-sheng Lin, Hui-lian Zhu, Hongwei Zhou, Yu-ming Chen, Ju-Sheng Zheng

## Abstract

Gut microbiome targets for type 2 diabetes (T2D) prevention among human cohorts have been controversial. Using an interpretable machine learning-based analytic framework, we identified robust human gut microbiome features, with their optimal threshold, in predicting T2D. Based on the results, we constructed a microbiome risk score (MRS), which was consistently associated with T2D across 3 independent Chinese cohorts involving 9111 participants (926 T2D cases). The MRS could also predict future glucose increment, and was correlated with a variety of gut microbiota-derived blood metabolites. Faecal microbiota transplantation from humans to germ-free mice demonstrated a causal role of the identified combination of microbes in the T2D development. We further identified adiposity and dietary factors which could prospectively modulate the MRS, and found that body fat distribution may be the key factor modulating the gut microbiome-T2D relationship. Taken together, we proposed a new analytical framework for the investigation of microbiome-disease relationship. The identified microbiota may serve as potential drug targets for T2D in future.

## Introduction

Type 2 diabetes (T2D) is a complex disorder influenced by both host genetic and environmental factors (*1*), and its prevalence is rising rapidly in both developed and developing countries (*2*). Gut microbiome is considered as a modifiable environmental factor, which plays an important role in the development of T2D (*3*–*7*). The research interest to identify gut microbiome-related treatment/prevention target is emerging recently (*8*). Although there are a few human studies investigating the association of gut microbiome with T2D in the past few years, the results are inconsistent, and the causality is lacking (*9*). So far, there are sparse human evidence robustly linking specific gut microbiome features to T2D.

Machine learning has been widely used in biomedical fields in recent years (*10*). However, its application in the clinical setting is still limited as their predictions are usually difficult to interpret. Of note, with the methodology development in the past few years, interpretable algorithms could unlock the traditional “black box” of machine learning results (*11*). The integration of the new algorithms with large-scale gut microbiome data have the potential to radically unveil the relationship between gut microbiome and T2D. Yet, no such investigation has been done.

Therefore, in the present study, we aimed to identify robust human gut microbiome features in predicting T2D with a novel interpretable machine learning analytical framework in large-scale human cohort studies. We also assessed the correlation between the combination of microbes and host blood metabolites to provide insight into the role of T2D-related gut microbiota in host metabolism. We further performed a faecal microbiota transfer experiment to establish the causality of the identified combination of microbes on the T2D development. As a secondary objective, we aimed to identify potential adiposity, dietary and lifestyle factors which could modify the T2D-related gut microbiota using our longitudinal cohort data.

## Results

### Linking host multi-dimensional information and T2D based on a machine learning method

The characteristics of the participants for the current study are shown in Table 1, and the overview of the study workflow is shown in Fig.1 and Fig.S1. 297 host features (metadata, gut microbiota composition, and gut microbiota diversity, see Supplemental text) were incorporated into our analyses. The metadata were collected at the same point-in-time as the stool sample. Prevalent T2D cases were ascertained on the basis of fasting blood glucose ≥7.0 mmol/L or HbA1c ≥6.5% or currently under medical treatment for diabetes at either of the follow-up visits, according to the American Diabetes Association criteria for the diagnosis of diabetes (*12*). We used LightGBM (*13*), a Gradient Boosting Decision Tree (GBDT) algorithm, to infer the relationship between incorporated features and T2D (Materials and Methods). Our machine learning model showed a high and robust performance for the prediction of T2D (AUC=0.86~0.89) in the discovery and external validation cohort 1 (Fig.2A, and Table S1). The LightGBM algorithm used in the present study outperformed the random forest algorithm in the T2D prediction (Table S2).

**Table 1.**
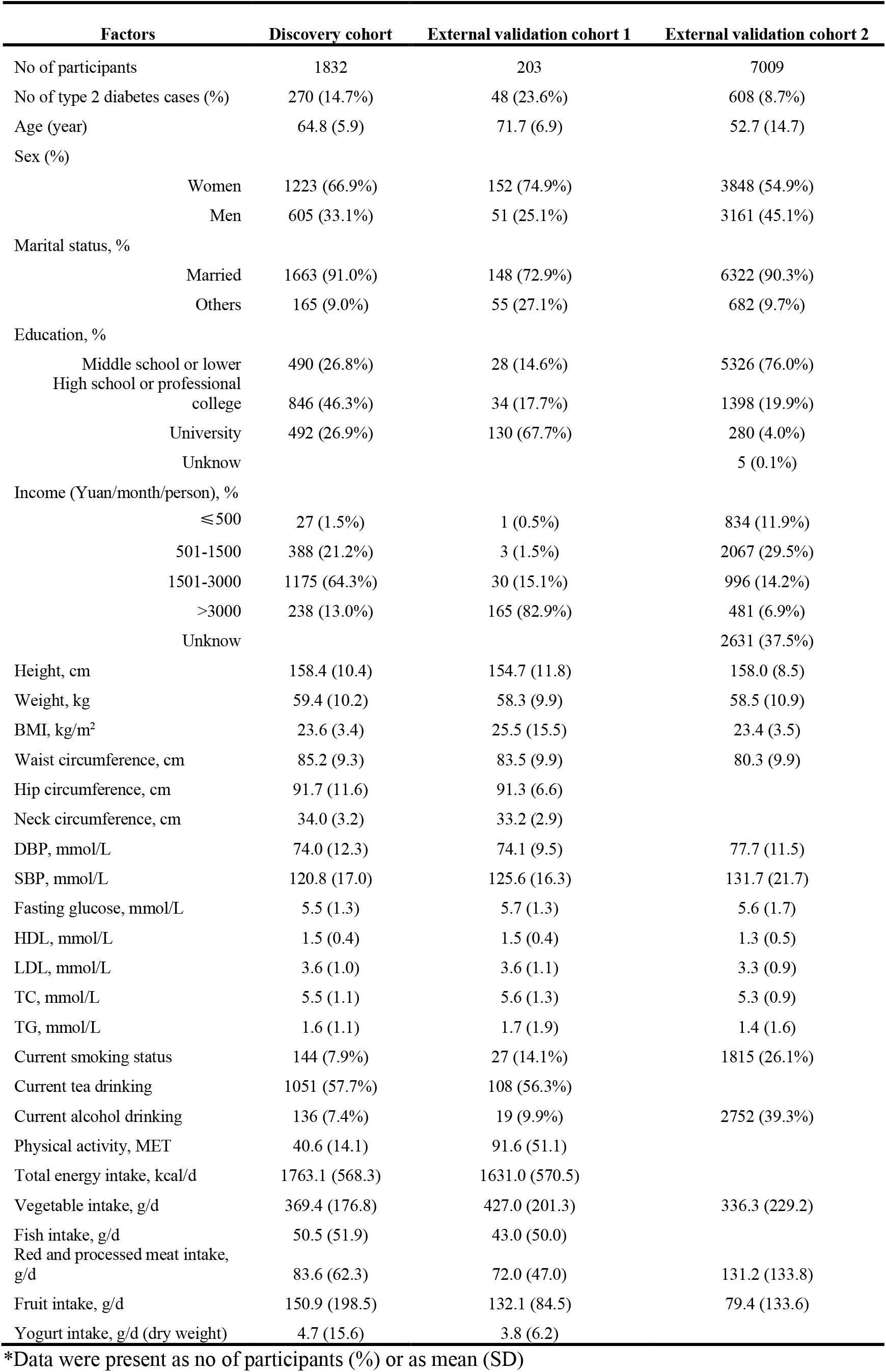
Characteristics of the participants included in this study*.

| Factors | Discovery cohort | External validation cohort 1 | External validation cohort 2 |
| --- | --- | --- | --- |
| No of participants | 1832 | 203 | 7009 |
| No of type 2 diabetes cases (%) | 270 (14.7%) | 48 (23.6%) | 608 (8.7%) |
| Age (year) | 64.8 (5.9) | 71.7 (6.9) | 52.7 (14.7) |
| Sex (%) |  |  |  |
| Women | 1223 (66.9%) | 152 (74.9%) | 3848 (54.9%) |
| Men | 605 (33.1%) | 51 (25.1%) | 3161 (45.1%) |
| Marital status, % |  |  |  |
| Married | 1663 (91.0%) | 148 (72.9%) | 6322 (90.3%) |
| Others | 165 (9.0%) | 55 (27.1%) | 682 (9.7%) |
| Education, % |  |  |  |
| Middle school or lower | 490 (26.8%) | 28 (14.6%) | 5326 (76.0%) |
| High school or professional college | 846 (46.3%) | 34 (17.7%) | 1398 (19.9%) |
| University | 492 (26.9%) | 130 (67.7%) | 280 (4.0%) |
| Unknow |  |  | 5 (0.1%) |
| Income (Yuan/month/person), % |  |  |  |
| ≤500 | 27 (1.5%) | 1 (0.5%) | 834 (11.9%) |
| 501-1500 | 388 (21.2%) | 3 (1.5%) | 2067 (29.5%) |
| 1501-3000 | 1175 (64.3%) | 30 (15.1%) | 996 (14.2%) |
| >3000 | 238 (13.0%) | 165 (82.9%) | 481 (6.9%) |
| Unknow |  |  | 2631 (37.5%) |
| Height, cm | 158.4 (10.4) | 154.7 (11.8) | 158.0 (8.5) |
| Weight, kg | 59.4 (10.2) | 58.3 (9.9) | 58.5 (10.9) |
| BMI, kg/m<sup>2</sup> | 23.6 (3.4) | 25.5 (15.5) | 23.4 (3.5) |
| Waist circumference, cm | 85.2 (9.3) | 83.5 (9.9) | 80.3 (9.9) |
| Hip circumference, cm | 91.7 (11.6) | 91.3 (6.6) |  |
| Neck circumference, cm | 34.0 (3.2) | 33.2 (2.9) |  |
| DBP, mmol/L | 74.0 (12.3) | 74.1 (9.5) | 77.7 (11.5) |
| SBP, mmol/L | 120.8 (17.0) | 125.6 (16.3) | 131.7 (21.7) |
| Fasting glucose, mmol/L | 5.5 (1.3) | 5.7 (1.3) | 5.6 (1.7) |
| HDL, mmol/L | 1.5 (0.4) | 1.5 (0.4) | 1.3 (0.5) |
| LDL, mmol/L | 3.6 (1.0) | 3.6 (1.1) | 3.3 (0.9) |
| TC, mmol/L | 5.5 (1.1) | 5.6 (1.3) | 5.3 (0.9) |
| TG, mmol/L | 1.6 (1.1) | 1.7 (1.9) | 1.4 (1.6) |
| Current smoking status | 144 (7.9%) | 27 (14.1%) | 1815 (26.1%) |
| Current tea drinking | 1051 (57.7%) | 108 (56.3%) |  |
| Current alcohol drinking | 136 (7.4%) | 19 (9.9%) | 2752 (39.3%) |
| Physical activity, MET | 40.6 (14.1) | 91.6 (51.1) |  |
| Total energy intake, kcal/d | 1763.1 (568.3) | 1631.0 (570.5) |  |
| Vegetable intake, g/d | 369.4 (176.8) | 427.0 (201.3) | 336.3 (229.2) |
| Fish intake, g/d | 50.5 (51.9) | 43.0 (50.0) |  |
| Red and processed meat intake, g/d | 83.6 (62.3) | 72.0 (47.0) | 131.2 (133.8) |
| Fruit intake, g/d | 150.9 (198.5) | 132.1 (84.5) | 79.4 (133.6) |
| Yogurt intake, g/d (dry weight) | 4.7 (15.6) | 3.8 (6.2) |  |

\*Data were present as no of participants (%) or as mean (SD)

**Fig.1.**
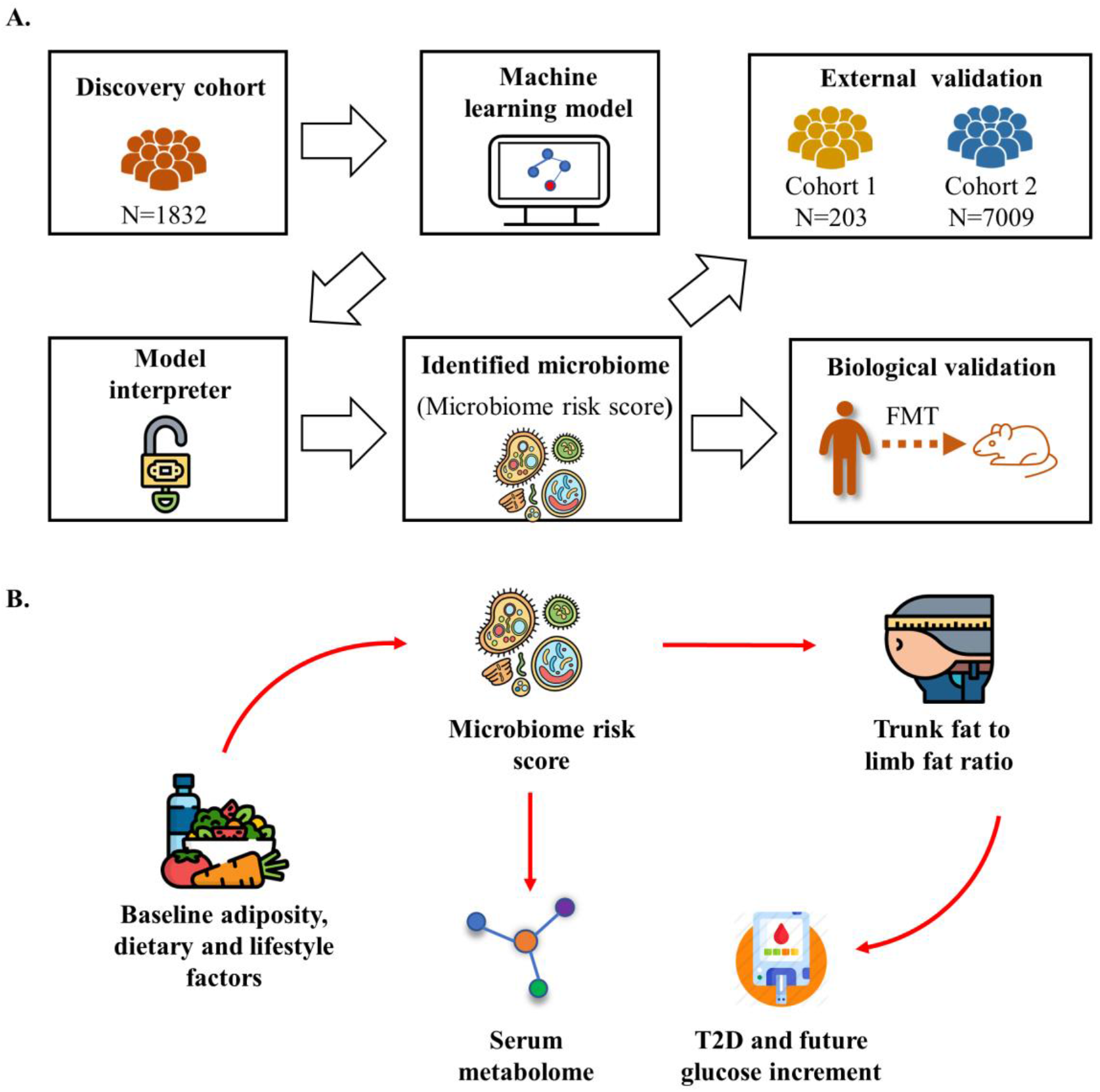
Study overview. **(A)** Identifying microbiome features, together with their optimal threshold and direction associated with type 2 diabetes (T2D). 1) Training and optimizing a machine-learning model to link the input factors with T2D in a discovery cohort (n=1832, 270 cases); 2) Using SHAP method to explain the output of machine learning model and identify the microbiome features associated with T2D risk; 3) Constructing a microbiome risk score (MRS) for T2D integrating the threshold and direction of the above-identified microbiome features. 4) Validating the MRS-T2D association in two independent external validation cohorts: cohort 1 (n=203, 48 cases), cohort 2 (n=7009, 608 cases); 5) Demonstrating a causal role of the identified microbiome in the T2D development by faecal microbiota transplantation (FMT). **(B)** Investigating the prospective association of baseline adiposity, dietary and lifestyle factors with the identified T2D-related microbiome features (i.e., MRS), and the correlation of the MRS with host serum metabolome. Further, we investigated the role of body fat distribution linking the MRS and T2D development in the discovery cohort and external validation cohort 1.

**Fig.2.**
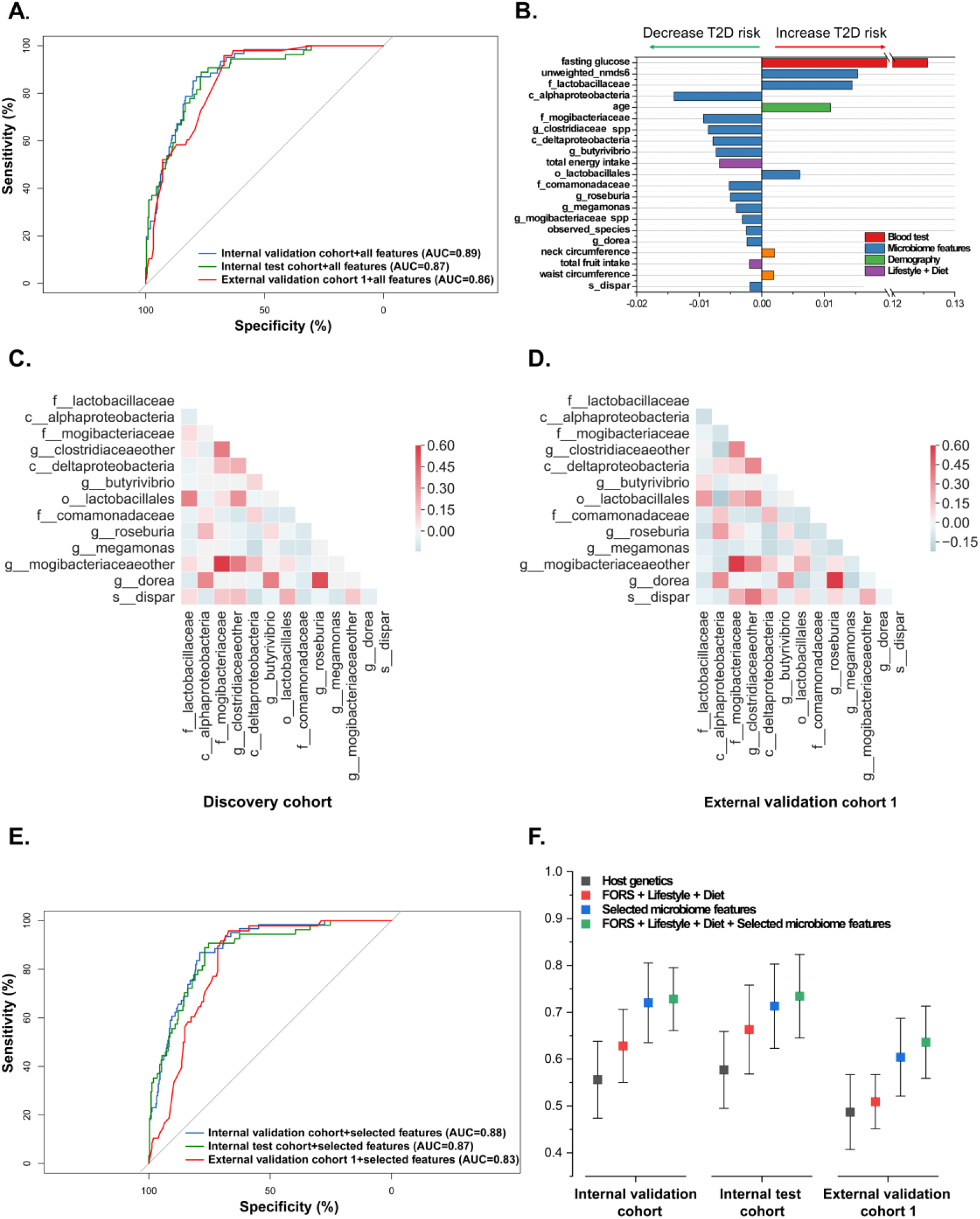
Linking host multi-dimensional information and type 2 diabetes (T2D) based on an interpretable machine learning framework. **(A)** Receiver Operator Characteristic curves (ROC curves) of the predictive models based on all 297 input features in the discovery cohort and external validation cohort 1. **(B)** The average impact of selected features on T2D risk. The bars are colored according to data categories. **(C-D)** The inter-correlation of selected microbiome features in the discovery cohort and external validation cohort 1. **(E)** ROC curves of the predictive models based on the selected features (n=21) in the discovery cohort and external validation cohort 1. **(F)** Algorithm performance in the discovery cohort and external validation cohort 1 based on the selected microbiome features, host genetics, lifestyle and diet, T2D traditional risk factors (FORS), and their combination.

### Factors underlying T2D prediction

To gain insight into the contribution of the different features in the algorithm’s prediction, we used SHapley Additive explanation(SHAP) (*11*) to interpret the machine learning model. Features with an average absolute SHAP value greater than 0 were used as selected features. We finally identified 21 features associated with the risk of T2D, of which 15 were microbiome features (two of them are indicators of microbial diversity, others are taxa-related features) (Fig.2B, Fig.S2 and Table S3), and the majority of the selected microbiome features had a low to modest intercorrelation (Fig.2, C to D, and Table S4). The selected features from the model showed a similar predictive capacity compared to all input features (Fig.2E, and Table S1).

We explored the marginal effect of each selected feature on T2D risk accounting for other features to examine how a single selected feature affected the output of the machine learning model. We created a SHAP dependence plot to show the effect of a single feature across the whole dataset (Fig.S3). Our results indicated that individuals with age >66.7 years or waist circumference >84.6cm were considered at high risk of T2D (Fig.S3). This is consistent with the standards of medical care for T2D in China (*14*, *15*), which suggests that individuals >65 years old or with waist circumference >85cm (male) or 80cm (female) are at high risk of T2D. These results further demonstrated the validity of our novel machine learning-based analytic framework.

We identified the optimal threshold of the identified 13 taxa-related features according to their SHAP dependence plots (Table S5). 8 of 13 taxa-related features showed statistically significant associations with T2D when they were treated as binary variables: high abundance (i.e., ≥ the optimal threshold) compared to low abundance (i.e., < the optimal threshold) (Fig.S4, A, and Table S6), while only 3 taxa-related features showed significant association with T2D if the abundance of the selected microbiome was treated as a continuous variable (Fig.S4, B, and Table S6). These results highlight the importance of our interpretable machine learning framework to identify the optimal threshold for the individual microbes, suggesting that a linear model may not be suitable for microbiome analysis.

### The identified combination of microbes is strongly predictive of T2D risk

To estimate individual microbiome risk for T2D development, we generated a microbiome risk score (MRS) integrating the threshold and direction of the above-identified microbial features (13 taxa-related features and observed species) to predict T2D risk (Materials and Methods). The MRS (ranges from 0-14) showed superior T2D prediction accuracy compared to the host genetics (T2D genetic risk score), Framingham-Offspring Risk Score (FORS) components (age, sex, parental history of diabetes, BMI, systolic blood pressure, high-density lipoprotein cholesterol, triglycerides, and waist circumference), lifestyle and dietary factors (current smoking status, current tea-drinking, current alcohol drinking, physical activity, total energy intake, vegetable intake, fish intake, red and processed meat intake, fruit intake and yogurt intake) (Fig.2F, and Table S7). An addition of the MRS to the model (FORS + lifestyle + diet) increased the AUC from 0.63 (95% CI 0.55-0.71) to 0.73 (95% CI 0.66-0.8) in the internal validation cohort (*P*=0.0024), 0.66 (95% CI 0.57-0.76) to 0.73 (95% CI 0.65-0.82) in the internal test cohort (*P*=0.016), and 0.51 (95% CI 0.45-0.57) to 0.64 (95% CI 0.56-0.71) in the external validation cohort 1 (*P*=0.0036), respectively.

We found that the MRS (per unit change in MRS) consistently showed positive association with T2D risk in the discovery cohort (RR 1.28, 95%CI 1.23-1.33), external validation cohort 1 (RR 1.23, 95%CI 1.13-1.34) and external validation cohort 2 (RR 1.12, 95%CI 1.06-1.18) (Fig.3A, Table 2, and Table S8). We also repeated the MRS-T2D association based on 1068 deep shotgun metagenomics samples in the discovery cohort (including 159 T2D cases). In agreement with the 16S RNA results, the metagenome-based MRS consistently showed positive association with T2D risk (per unit change in new MRS: RR 1.33, 95%CI 1.17-1.51) (Fig.3A, and Table S8).

**Fig.3.**
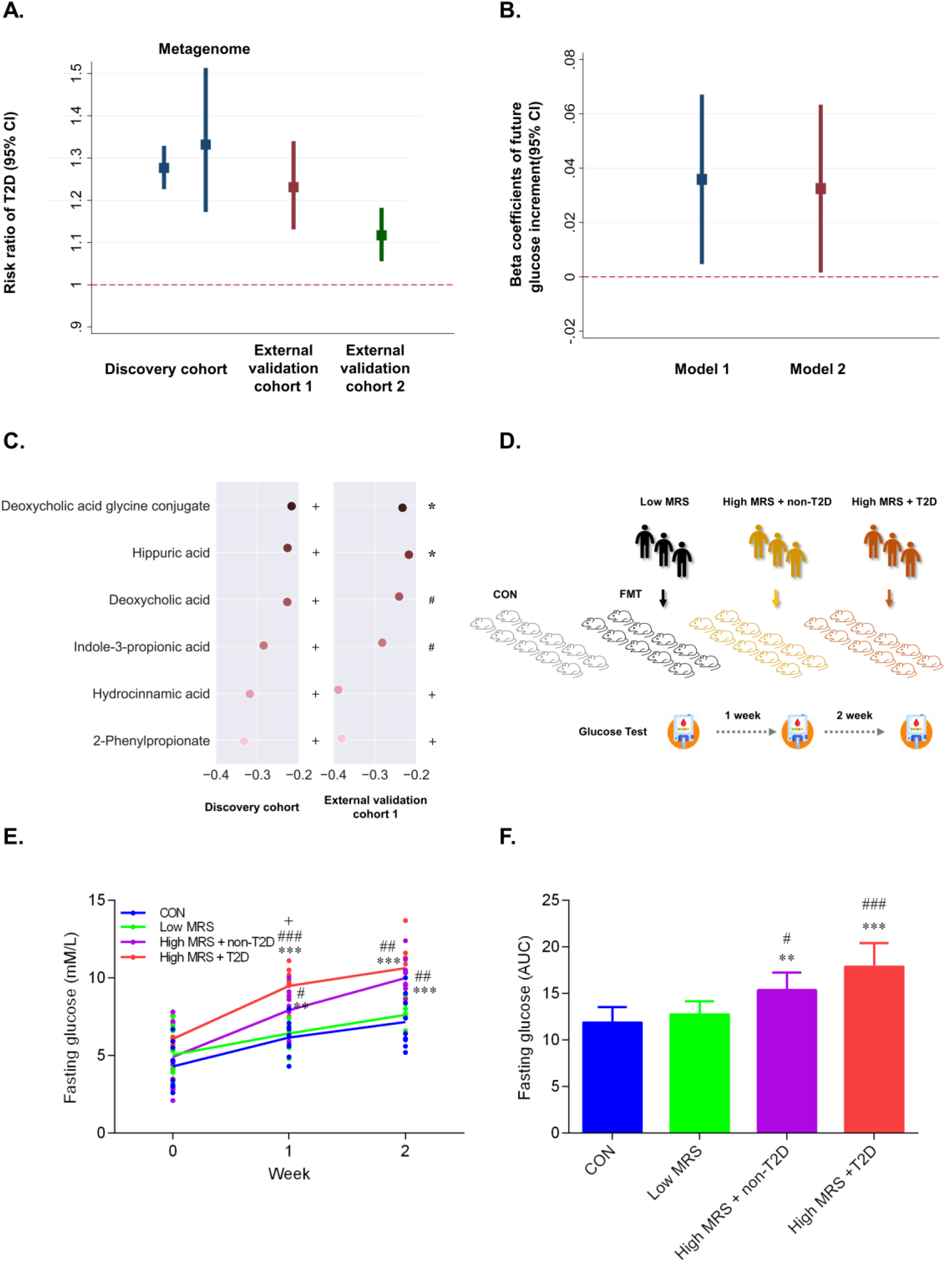
Identified gut microbiota affect the type 2 diabetes (T2D) development and host serum metabolites. **(A)** Association of the microbiome risk score (MRS) with T2D risk in discovery cohorts, external validation cohort 1, and external validation cohort 2. Poisson regression was used to estimate the risk ratio (RR) and 95% confidence interval (CI) of T2D per unit change in the MRS, adjusting for demographic, dietary and lifestyle factors. **(B)** Association between the MRS and prospective glucose increments over 3 years in discovery cohort. Linear regression was used to estimate the difference in future fasting glucose per unit change in the MRS in a cohort of 249 non-T2D individuals, adjusted for demographic, dietary and lifestyle factors (model 1). Sensitivity analyses were conducted under model 1 by plus baseline fasting glucose to test the influence of baseline fasting glucose on the performance of our model (model 2). **(C)** Association of the microbiome risk score (MRS) with host circulating metabolites. The Spearman correlation coefficients between the microbiome risk score and the host serum metabolites were calculated. The MRS-metabolite associations were further replicated in the external validation cohort 1. * *P*< 0.05, # *P*< 0.01, + *P*< 0.001. **(D-F)** Identified gut microbiota causally affect the type 2 diabetes (T2D) development in germ-free mice. **(D)** Schematic diagram. **(E)** Fasting glucose curves. **(F)** Quantification of fasting glucose by AUC. * compared with CON group, # compared with Low MRS group, + compared with High MRS+non-T2D group. (*, #, +) *P*< 0.05, (**, ##, ++) *P*< 0.01, (***, ###, +++) *P*< 0.001 by ANOVA. The *P*-values were adjusted using the Benjamini and Hochberg method.

**Table 2.** Association of the gut microbiome risk score (MRS) with type 2 diabetes*.

| Cohorts | Median (MRS) | No. of cases / Total No. | Adjusted risk ratio (95% CI) | P value |
| --- | --- | --- | --- | --- |
| **Discovery cohort** |  |  |  |  |
| Q1 | 3 | 33 / 569 | 1 (reference) |  |
| Q2 | 5 | 62 / 515 | 2.02 (1.35, 3.02) | <0.001 |
| Q3 | 7 | 70 / 419 | 2.73 (1.85, 4.04) | <0.001 |
| Q4 | 10 | 101 / 304 | 5.29 (3.66, 7.65) | <0.001 |
| **External validation cohort 1** |  |  |  |  |
| Q1 | 4 | 7 / 65 | 1 (reference) |  |
| Q2 | 6 | 4 / 31 | 1.47 (0.49, 4.43) | 0.49 |
| Q3 | 7 | 15 / 53 | 2.6 (1.17, 5.79) | 0.019 |
| Q4 | 10 | 17 / 39 | 4.17 (1.96, 8.85) | <0.001 |
| **External validation cohort 2** |  |  |  |  |
| Q1 | 6 | 236 / 3065 | 1 (reference) |  |
| Q2 | 7 | 147 / 1672 | 1.11 (0.91, 1.35) | 0.31 |
| Q3 | 8 | 110 / 1104 | 1.27 (1.03, 1.57) | 0.025 |
| Q4 | 9 | 104 / 946 | 1.36 (1.10, 1.68) | 0.0051 |

\*Poisson regression was used to estimate the risk ratio (RR) and 95% confidence interval (CI) of the type 2 diabetes in each of the three cohorts, according to the gut microbiome risk score. In these comparisons, participants at low microbiome risk (Q1) were treated as the reference group. The covariates for the discovery cohort and validation cohort 1 were total energy intake, age, waist circumference, sex, BMI, alcohol status, smoking status, education, marital status and income. For the validation cohort 2 (GGMP), covariates including age, waist circumference, sex, BMI, alcohol status, smoking status, education, marital status.

### The identified combination of microbes is longitudinally related with glucose increments

In order to investigate the relationship between the identified combination of microbes (i.e., MRS) and glucose increments longitudinally. We conducted a prospective investigation among 249 GNHS cohort participants with normal fasting glucose (fasting glucose <7 mmol/l) at baseline, who were followed up for a median of 3.4 years after the collection of stool samples. Linear regression was used to calculate the correlation coefficient (Beta) and 95% confidence interval (CI) of glucose increments per unit higher in the MRS after adjusting for age, sex, BMI, waist circumference, smoking status, household income, alcohol drinking status, total energy intake, marital status and education level (model 1). We also conducted a sensitivity analysis to test the influence of baseline fasting glucose on the performance of our model by including baseline fasting glucose into the model. Our results showed that MRS was significantly positively associated (*P*<0.05) with future glucose increments in two statistical models (Fig.3B, and Table S9). These results indicate that our identified combination of microbes could predict future glucose status among non-T2D participants.

### Correlation of the identified combination of microbes with host blood metabolome

We performed targeted metabolomics profiling of serum samples from the discovery cohort (n=903) and external validation 1 (n=113), and assessed the correlation of the T2D-related combination of microbes (i.e., MRS) with 199 serum metabolites (Supplemental text). Participants with a history of the T2D medication use were excluded in this analysis. The serum samples were collected at the same point-in-time as the stool samples. We found the MRS was consistently correlated with 6 metabolites in the discovery cohort and external validation cohort 1 (Fig.3C).

The MRS was negatively correlated with 2-phenylpropionate, hydrocinnamic acid and indole-3-propionic acid, which were all associated with gut microbiome metabolism (*16*–*18*). Deoxycholic acid and deoxycholic acid glycine conjugate are secondary bile acids produced by the action of enzymes existing in the microbial flora of the colonic environment (*19*). Recent studies have revealed that alteration of gut microbiota could not only affect the bile acid pool, but also influence the bile acid receptor signaling (i.e., FXR and TGR5). The FXR has been reported to be involved in glucose homeostasis, energy expenditure, and lipid metabolism (*20*).These observations provide insight into the potential function and mechanism of our identified microbial features, represented by the MRS, in host metabolism.

### The identified combination of microbes causally affect the T2D development in germ-free mice

To determine the causality between the identified combination of microbes and T2D risk, we transferred human faecal samples to germ-free mice to investigate the effects of the identified microbiota on T2D development (Fig.3D, Materials and Methods). Mice transplanted with the gut microbiota from high MRS individuals, either at non-T2D or T2D status, showed significant increase in fasting glucose levels compared with those from the low MRS individuals or germ-free control mice (Fig.3E to F). There was no significant difference in fasting glucose between the germ-free control group and the low MRS group. The mice weight of each group during follow-up was shown in Fig.S5 A to B. These results provide evidence for a causal relationship of the selected gut microbial features with T2D risk.

### Baseline adiposity and dietary factors can modulate the T2D-related microbiome

We examined whether the MRS could be modulated by baseline adiposity, dietary or lifestyle factors (components see table S10). In the longitudinal analysis of the discovery cohort, baseline BMI were positively associated with the MRS, while hip circumference and tea-drinking was inversely associated (Fig.4A, and Table S10).

**Fig.4.**
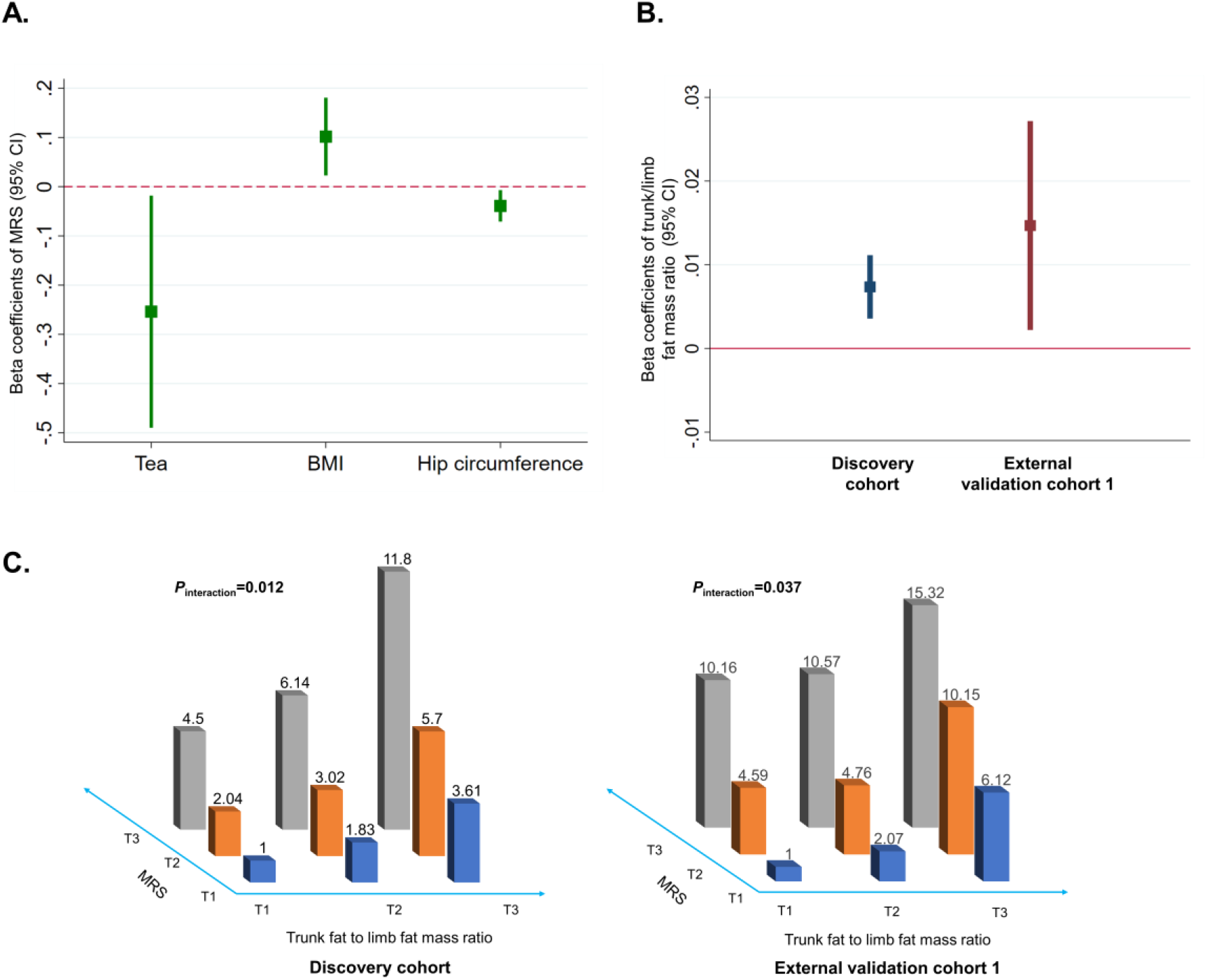
Adiposity and dietary factors modulate the association between gut microbiome and type 2 diabetes (T2D). **(A)** Association of baseline adiposity and dietary factors with the microbiome risk score (MRS). Linear regression was used to estimate the difference in MRS per quartile (for continuous dietary factors) or per unit (for adiposity factors) or per category (for ordinary factors) change in the baseline tested factors, adjusted for demographic factors, T2D medication use, and mutually adjusted for the other tested adiposity, dietary and lifestyle factors. We only presented those adiposity, dietary or lifestyle factors showing significant association with the MRS in the figure. **(B)** Association between the MRS and trunk fat to limb fat mass ratio in discovery cohort and external validation cohort 1. Linear regression was used to estimate the difference in trunk fat to limb fat mass ratio per unit change in the MRS, adjusted for demographic, dietary and lifestyle factors. **(C)** Interaction between MRS and trunk fat to limb fat mass ratio on T2D risk. Poisson regression was used to estimate the interaction of MRS and trunk fat to limb fat mass ratio on T2D risk, adjusted for demographic, dietary and lifestyle factors

### Body shape is associated with gut microbiome, modulating the association of gut microbiome with T2D

Obesity is a most important risk factor of T2D (*21*). As BMI and hip circumference are closely correlated with the MRS in our study, we hypothesized that the relationship of gut microbiome with T2D might be modulated by the adiposity status. The MRS was positively associated (*P*<0.05) with the distribution of trunk to limb fat ratio (trunk/limb fat mass ratio) in the discovery cohort and external validation cohort 1 (Fig.4B and Table S11-Table S12). We found a significant interaction between MRS and trunk/limb fat mass ratio for T2D risk in the discovery cohort (*P*_interaction_=0.012) and external validation cohort1 (*P*_interaction_=0.037), adjusted for potential confounders (Fig.4E). In the discovery cohort, adjusted risk ratio (95% CIs) of T2D according to tertiles of the trunk/limb fat mass ratio was 1 (reference), 1.83 (0.86-3.88) and 3.61 (1.81-7.18) in the lowest MRS tertile, and 4.5 (2.21-9.17), 6.14 (3.12-12.08) and 11.79 (6.28-22.16) in the highest MRS tertile. Similar interaction results were found in the external validation cohort 1 (Fig.4C, and Table S13).

## Discussion

In the present study we identify robust combination of microbes in predicting T2D by integrating a cutting-edge interpretable machine learning framework with large-scale human cohort studies. We construct a novel risk score for the gut microbiome, which shows superior T2D prediction accuracy compared to host genetics or traditional risk factors. Additionally, we successfully replicate the MRS-T2D association in another two independent cohorts. We then reveal that the MRS is correlated with a few gut microbiota-derived blood metabolites. The faecal microbiota transfer experiment confirmed the causality of the identified combination of microbes on T2D development. Finally, we identify potential baseline factors which could modulate the T2D-related microbiome features, and demonstrate that the relationship between the microbiome and T2D could be modified by the body fat distribution.

Microbiome data are highly dimensional, underdetermined, over-dispersed, and often sparse with excess zeros. These features challenge standard statistical tools, making results from both traditional parametric and non-parametric models unsatisfactory (*22*). On the other hand, multiple host anthropometric, dietary and lifestyle factors play important roles in shaping the microbiome composition (*23*–*25*); while large human cohorts that taking into account these confounders are necessary but are so far sparse. The machine learning algorithm (LightGBM) we used to integrate host demographic, clinical, dietary, lifestyle and microbiome profiles outperformed the random forest algorithm in the T2D prediction. We also interpret the results of the ‘black box’ machine learning models with a recently developed novel tool: SHAP (*11*). Compared with other interpreting methods such as gain, split count and permutation method, SHAP has been theoretically verified as the only consistent and locally accurate method to interpret machine learning results (*26*). We demonstrated that our new analytic framework could effectively integrate data from different dimensions and subsequently unlocking the machine learning-generated ‘black box’ results. This analytic framework could be used for other multi-omics research as well, beyond gut microbiome.

The first published human cohort study examining the difference of gut microbiome between T2D cases (n=18) and healthy controls (n=18) found that proportions of phylum *Firmicutes* and class *Clostridia* were significantly reduced in the T2D group compared to the control group (*5*). However, these results were not confirmed in another two small human gut microbiome studies conducted in China and Europe (*27*, *28*). Although results from the above two studies (*27*, *28*) suggested that functional alterations of the gut microbiome might be directly linked to T2D development, the most discriminatory microbial markers for T2D differ between the two studies.

Most of our identified T2D-related taxa were from the order *Clostridiales* (*f_mogibacteriaceae, g_clostridiaceae spp, g_butyrivibrio, g_roseburia, g_megamonas, g_mogibacteriaceae spp, g_dorea, s_dispar*), which were consistently enriched in the healthy controls, rather than T2D cases. Specifically, *roseburia*, which is decreased in our T2D patients, is a butyrate-producing genus and has been shown to causally improve glucose tolerance (*29*, *30*). A previous study has demonstrated that reduction in the diversity and function of the class *Clostridia* contributes to the obesity development potentially via down-regulated genes that control lipid absorption (*31*). Therefore, the potential effect of *Clostridia* on obesity may explain our observed interaction between MRS and body fat distribution. In line with previous literature indicating that genus *lactobacillus* might contribute to chronic inflammation in diabetes development (*5*, *32*), we also found that the family *lactobacillaceae* was enriched in the T2D participants and had a strong predictive power for T2D. Although based on the different microbiome analysis method, the two shotgun metagenomics based studies (*27*, *28*) consistently showed a decrease in *roseburia* species and an increase in *lactobacillus* species in T2D cases compared to controls. Specially, *lactobacillus* species had the highest score for the identification of T2D patients in a European study (*28*). Due to the translational nature of the present project, we did not further investigate the functionality of each identified gut microbial taxa, but rather, we were more interested in the role of the overall microbiome combination and pattern.

We developed the concept of MRS for T2D. The MRS could predict future glucose change prospectively, inferring the potential causality of the identified combination of microbes in diabetes development, which was confirmed by our faecal microbiota transplantation study. The prospective investigation of the gut microbiome-glucose association was rarely conducted by any of the previous cohort studies, which exclusively investigated a cross-sectional association of gut microbiome with T2D or related traits (*5*, 9, *27*, *28*, *33*–*35*). Integration of MRS-blood metabolome analysis revealed potential mechanism of the MRS-T2D association, involving a variety of gut microbiota-derived metabolites, although the detailed mechanism is yet to be discovered.

We further demonstrated that higher BMI or lower hip circumference is positively associated with future MRS levels, which indicates the potential role of adiposity in affecting gut microbiome. The evidence is clearer when we found an interaction between the MRS and trunk to limb fat mass ratio, suggesting that adiposity may be an effect modifier for gut microbiome and T2D development. Taken together, our results highlight that a healthy body shape may play an important role in maintaining the gut health.

In summary, with a high-accuracy machine learning model and a credible interpreter, we discover and validate the associations of gut microbiome and the related MRS with T2D in several large human cohorts. These newly discovered combination of microbes can be potentially used as T2D diagnostic, therapeutic targets, or preventive targets through diet and lifestyle intervention. Furthermore, the MRS can potentially assist in the screening of the best faecal donors for the treatment of T2D patients in future and improve the clinical therapeutic safety of faecal transplantation.

## Materials and Methods

### Study design

We included participants from three human cohorts, 1832 participants from the Guangzhou Nutrition and Health Study (GNHS) (*36*) as a discovery cohort (270 T2D cases), 203 participants belonged to the control arm of a case-control study of hip fracture in Guangdong Province, China (*37*) as an external validation cohort 1 (48 T2D cases), and another 7009 participants from GGMP (Guangdong Gut Microbiome Project) as an further external validation cohort 2 (608 cases) (*23*). Detailed study designs of GNHS have been reported previously(*36*). Briefly, GNHS is an ongoing community-based prospective cohort study in Guangzhou, China. There were two waves of participant recruitment using the same criteria: between 2008 and 2010 (n=3169), and between 2012 and 2013 (n=879). All participants were followed up every 3 years. Stool samples were collected at the second and third follow-up. Those with measurement of 16s rRNA from stool samples were included in the present study (n=1935). Study participants were excluded if they had an unclear diabetes status (n=48), chronic renal dysfunction or self-reported cancers (n=55). Finally, 1832 participants were included in the present analysis as a discovery cohort, including 1068 individuals (159 T2D cases) with a measurement of shotgun metagenomic sequence. Among the included participants, there were 249 non-T2D participants, who were followed up for a median of 3.4 years after the collection of their stool samples. These participants were included in our longitudinal analysis of gut microbiome with glucose increments. All 1832 participants were included in our longitudinal analysis on the prospective association of baseline factors with gut microbiome (with a median follow up of 6.2 years).

The hip fracture case-control cohort (external validation cohort 1) was performed between June 2009 and June 2012 in Guangdong Province, China. Detailed information of this cohort has been reported previously (*37*). After adopting the same inclusion and exclusion criteria as GNHS, we included 203 participants with a measurement of 16s rRNA from stool samples in the present analysis. The study protocols of GNHS and the hip fracture case-control study were approved by the Ethics Committee of the School of Public Health at Sun Yat-sen University, and all participants gave written informed consent.

Details method for the covariate measurements, stool sample collection, 16s rRNA sequencing, shotgun metagenome sequencing and taxonomy analysis for GNHS and hip fracture case-control study was provided in Supplemental text.

All GGMP participants (external validation cohort 2) were from 14 randomly selected districts or counties in Guangdong province. In each district or county, three neighborhoods or townships were selected, and in each neighborhood or township, two communities or villages were selected (*23*). Detailed methods for the assessment of demographic, lifestyle and dietary information, stool sample collection, processing and 16s sequencing for GGMP have been reported previously (*23*). The study protocol was approved by the Ethical Review Committee of the Chinese Center for Disease and Prevention, and all participants gave written informed consent.

### Interpretable machine learning framework for data integration and explanation

We devised a model based on a gradient boosting framework —LightGBM(*13*) to link input features with T2D (detailed parameters were provided in Supplementary text).

To train and validate our model, we divided the discovery cohort into three parts randomly at a ratio of 6:2:2, resulting in 1099, 366 and 367 participants, which were allocated at the training cohort, internal validation cohort, and internal test cohort, respectively. The training cohort was used to fit parameters of the model; the internal validation cohort was used to tune parameters of the model; and the internal test cohort was used to assess the performance of the model. AUC was used to evaluate the model’s performance. Our predictor is based on code adapted from the sklearn 0.15.2 (*38*) lightgbm class, R packages pROC (*39*) were used for ROC curve analyses, “delong” method for AUC comparison. We also compared our model performance with that of a random forest algorithm, applying the same evaluation criteria (tenfold cross-validation in the discovery cohort, independent validation in the external cohort 1).

We used the SHAP (Shapley Additive exPlanations) (*11*) integrated into LightGBM to unlock the machine learning results. The inflection point of SHAP dependence plots (X-axis represents the feature variable, while Y-axis represents the SHAP value for the feature variable) were defined as the optimal threshold for each selected feature.

### Microbiome risk score (MRS) construction

We construct an MRS based on the machine learning-selected microbiome features and their SHAP values by using the additive model:

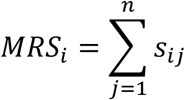

Where, *MRS*_i_ is a MRS for individual *i*, 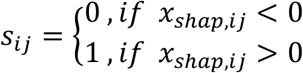, *s*_*ij*_is the microbiome risk score for the *jth* microbiome features in *ith* individual*. n* is the sum of the microbiome features, and *x*_*shap,ij*_ is the SHAP value for the *jth* microbiome features in *ith* individual. The MRS components including observe species, *f_lactobacillaceae, c_alphaproteobacteria, f_mogibacteriaceae, g_clostridiaceae spp, c_deltaproteobacteria, g_butyrivibrio, o_lactobacillales, f_comamonadaceae, g_roseburia, g_megamonas, g_mogibacteriaceae spp, g_dorea, s_dispar*.

### Gut microbiota transplantation

Nine participants were randomly selected as the representative donors according to the level of the MRS (ranges from 0-14):

1. Low MRS group: 3 participants, MRS=0, or MRS=1.
2. High MRS + non-T2D group: 3 participants, MRS=11.
3. High MRS + T2D group: 3 participants, MRS=13, or MRS=14.

Weaned, germ-free male C57BL/6J mice (*n* = 40) were maintained in flexible-film plastic isolators under a regular 12-h light cycle (lights on at 06:00). The mice were fed a sterilized normal chow diet (10% energy from fat; 3.25 kcal/g; SLAC). At 4 weeks of age, the germ-free mice were housed in individual cages and randomly divided into four groups (each group was kept in an individual isolator). After 1 weeks of acclimatization, the CON group of mice (*n* = 10) were orally gavaged with 100 μL of normal saline, and the other three groups of mice (*n* = 10, per group) were orally gavaged with 100 μL of the fecal suspension inoculum (taken from the each of the above donor group, preparation methods see supplementary materials). All mice were fed a sterilized high-fat diet. On Day 0, 7 and 14, after 12 h of fasting, fasting glucose was measured through the tail vein (Sinocare, China).

Detailed description of fecal suspension inoculum preparation was provided in Supplementary text. All animal experimental procedures were approved by the Ethics Committee of Westlake University and were conducted according to the committee’s guidelines.

### Statistical analysis

Statistical analysis was performed using Stata 15 (StataCorp, College Station, TX, USA). For the discovery cohort and external validation cohort 1, multivariable Poisson regression model (with robust standard errors) was used to examine the cross-sectional association with T2D for each machine-learning identified taxa-related feature as a continuous variable or as a binary variable: higher abundance (i.e., ≥the optimal threshold) compared with those lower abundance (i.e., <the optimal threshold), adjusted for age, sex, BMI, waist circumference, household income, marital status, and self-reported educational level, total energy intake, alcohol drinking, and smoking. For external validation cohort 2, all aforementioned covariates but total energy intake (not available) were used in the statistical model. We combined the effect estimates from the 3 cohorts using random-effects meta-analysis.

With the machine-learning identified MRS, in each of the internal validation cohort, internal test cohort and external validation cohort 1, we calculated the AUC for T2D prediction for the MRS, host genetics (T2D genetic risk score), and the traditional T2D risk factors including the Framingham-Offspring Risk Score (FORS) components (age, sex, parental history of diabetes, BMI, systolic blood pressure, high-density lipoprotein cholesterol, triglycerides, and waist circumference), lifestyle and dietary factors (current smoking status, current tea-drinking, current alcohol drinking, physical activity, total energy intake, vegetable intake, fish intake, red and processed meat intake, fruit intake and yogurt intake). ROC curves were compared with a paired two-sided DeLong’s test using the pROC package in R (*23*).

We also used a Poisson regression model (with robust standard errors) to explore the cross-sectional association of the MRS with T2D risk in our discovery cohort, and two external validation cohorts, respectively, adjusted for the same covariates as above individual taxa analysis. Given the information on household income was missing for many participants (n=2566, 37.8%) in external validation cohort 2, we performed sensitivity analysis by excluding household income as a covariate.

We used a linear regression model to explore the association of baseline MRS with glucose increments in the next 3 years, adjusted for the demographic and dietary and lifestyle factors. Sensitivity analysis was conducted by adding baseline fasting glucose to test the influence of baseline fasting glucose on the performance of the above model.

The association of the MRS with host circulating metabolites was assessed by the Spearman correlation. Those MRS-metabolite associations survived the multiple test correction (Benjamini and Hochberg method) in the discovery cohort were further chosen for replication in the external validation cohort 1.

In the discovery cohort, linear regression was used to estimate the difference in MRS per quartile change for continuous dietary factors or per unit change for adiposity factors or per category change for categorical (ordinary) factors in the baseline tested factors, adjusted for demographic factors, T2D medication use, and mutually adjusted for the other tested adiposity, dietary and lifestyle factors. The tested adiposity, dietary and lifestyle factors including BMI, hip circumference, waist circumference, neck circumference, total energy intake, alcohol drinking, smoking, tea drinking, vegetable intake, fruit intake, fish intake, red and processed meat intake, yogurt intake and physical activity. The adjusted demographic factors including age, sex, household income, marital status and educational level.

In both the discovery cohort and the external validation cohort 1, we used a linear regression model to assess the cross-sectional association of MRS with body fat distribution, adjusted for age, sex, total energy intake, alcohol drinking, smoking, household income, marital status and educational level. In both cohorts, Poisson regression was used to estimate the interaction of MRS with trunk fat to limb fat mass ratio on T2D risk, adjusted for the demographic, dietary and lifestyle factors.

For the results of the animal study, ANOVA was used for comparison between multiple groups. The *P*-values were adjusted using the Benjamini and Hochberg method. *P* values <0.05 were considered significant.

## Acknowledgments

We thank all of the participants of the cohorts for contributing stool samples and phenotypes.

## Author contributions

Conceptualization, J.S.Z., W.L.G., Y.M.C.; Methodology, J.S.Z and W.L.G.; Formal Analysis, W.L.G. and Z.L.J.; Investigation, C.W.L., Y.H., J.S.L., T.Y.S., and H.L.Z.; Data curation, C.W.L., Y.H., F.Z.X. and Z.L.M.; Resources, Y.M.C., H.W.Z. and J.S.Z.; Writing, W.L.G. and J.S.Z.; Writing-Review & Editing, J.S.Z., W.L.G., Y.Q.F., H.W.Z., Y.M.C., Y.H., Z.L.J, C.W.L., F.Z.X., Z.L.M., T.Y.S., J.S.L., and H.L.Z.; Visualization, W.L.G.; Supervision, J.S.Z., Y.M.C., and H.W.Z.; Funding Acquisition, J.S.Z., Y.M.C. and H.W.Z.

## Data and materials availability

For the discovery and external validation cohort 1, the raw data for 16 S rRNA gene sequences are available in the CNSA (https://db.cngb.org/cnsa/) of CNGBdb at accession number CNP0000829. For the external validation cohort 2, the raw data for 16 S rRNA gene sequences are available from the European Nucleotide Archive (https://www.ebi.ac.uk/ena/) at accession number PRJEB18535.

