## supplemental methods and tables for "Interpretable machine learning framework reveals novel gut microbiome features in predicting type 2 diabetes"

**Supplementary appendix**

**Supplemental text**

**Metadata in GNHS and the hip fracture case-control study**

Metadata included in this study was further categorized into 4 groups:

1) 5 demographic factors: age, sex, household income, marital status and self-reported educational level.

2) 10 lifestyle and dietary factors: physical activity, total energy intake, alcohol drinking, smoking, tea drinking, vegetable intake, fruit intake, fish intake, red and processed meat intake, and yogurt intake.

3) 5 blood test factors: Fasting glucose, HDL, LDL, TC, and TG.

4) 8 anthropometry factors: height, weight, hip circumference, waist circumference, neck circumference, BMI, DBP, SBP.

Description of each factor in different cohorts is listed in Table 1.

Demographic, lifestyle and dietary factors were all collected by questionnaire during on-site face-to-face interviews. Habitual dietary intakes over the past 12 months were assessed by a food frequency questionnaire, as previously described (*40*). Physical activity was assessed as a total metabolic equivalent for task (MET) hours per day on the basis of a validated questionnaire for physical activity (*41*). Anthropometric factors were measured by trained nurses on site during the baseline interview. Fasting venous blood samples were taken at each recruitment or follow-up visit. Serum low-density lipoprotein cholesterol and glucose were measured by coloimetric methods using a Roche Cobas 8000 c702 automated analyzer (Roche Diagnostics GmbH, Shanghai, China). Intra-assay coefficients of variation (CV) was 2.5% for glucose. Insulin was measured by electrochemiluminescence immunoassay (ECLIA) methods using a Roche cobas 8000 e602 automated analyzer (Roche Diagnostics GmbH, Shanghai, China). High-performance liquid chromatography was used to measure glycated hemoglobin (HbA1c) using the Bole D-10 Hemoglobin A1c Program on a Bole D-10 Hemoglobin Testing System, and the intraassay CV was 0.75%. The whole-body composition was measured by dual-energy x-ray absorptiometry (DXA) (Discovery W; Hologic Inc.). We analyzed the lean mass, fat mass and bon mass of the whole body, arms, and legs using the Hologic Discovery software version 3.2 (*42*).

**Stool sample collection and DNA extraction**

The stool samples were collected at a local study site within the School of Public Health at Sun Yat-sen University, and were transferred to a -80°C facility within 4 hours after collection. Total bacterial DNA was extracted using the QIAamp® DNA Stool Mini Kit (Qiagen, Hilden, Germany) following the manufacturer’s instructions. DNA concentrations were measured using the Qubit quantification system (Thermo Scientific, Wilmington, DE, US). The extracted DNA was then stored at -20 °C.

**16S gene amplicon sequencing**

The 16S rRNA gene amplification procedure was divided into two PCR steps, in the first PCR reaction, the V3-V4 hypervariable region of the 16S rRNA gene was amplified from genomic DNA using primers 341F(CCTACGGGNGGCWGCAG) and 805R(GACTACHVGGGTATCTAATCC). Amplification was performed in 96-well microtiter plates with a reaction mixture consisting of 1X KAPA HiFi Hot start Ready Mix, 0.1µM primer 341 F, 0.1 µM primer 805 R, and 12.5 ng template DNA giving a total volume of 50 µL per sample. Reactions were run in a T100 PCR thermocycle (BIO-RAD) according to the following cycling program: 3 min of denaturation at 94 °C, followed by 18 cycles of 30 s at 94 °C (denaturing), 30 s at 55 °C (annealing), and 30 s at 72 °C (elongation), with a final extension at 72 °C for 5 min. Subsequently, the amplified products were checked by 2% agarose gel electrophoresis and ethidium bromide staining. Amplicons were quantified using the Qubit quantification system (Thermo Scientific, Wilmington, DE, US) following the manufacturers’ instructions. Sequencing primers and adaptors were added to the amplicon products in the second PCR step as follows 2 µL of the diluted amplicons were mixed with a reaction solution consisting of 1×KAPA HiFi Hotstart ReadyMix, 0.5µM fusion forward and 0.5µM fusion reverse primer, 30 ng Meta-gDNA(total volume 50 µL). The PCR was run according to the cycling program above except with cycling number of 12. The amplification products were purified with Agencourt AMPure XP Beads (Beckman Coulter Genomics, MA, USA) according to the manufacturer’s instructions and quantified as described above. Equimolar amounts of the amplification products were pooled together in a single tube. The concentration of the pooled libraries was determined by the Qubit quantification system. Amplicon sequencing was performed on the Illumina MiSeq System (Illumina Inc., CA, USA). The MiSeq Reagent Kits v2 (Illumina Inc.) was used. Automated cluster generation and 2 × 250 bp paired-end sequencing with dual-index reads were performed.

**16S rRNA gene sequence data processing**

Fastq-files were demultiplexed by the MiSeq Controller Software (Illumina Inc.). The sequence was trimmed for amplification primers, diversity spacers, and sequencing adapters, merge-paired and quality filtered by USEARCH. UPARSE was used for OTU clustering equaling or above 97%. Taxonomy of the OTUs was assigned and sequences were aligned with RDP classifier. The OTUs were analyzed by phylogenetic and operational taxonomic unit (OTU) methods in the Quantitative Insights into Microbial Ecology (QIIME) software version 1.9.0 (*43*). α-diversity (Observed OTU number, Shannon index, Simpson index, Chao1 index, Goods coverage index) and β-diversity (Unweight UniFrac distances and Weight UniFrac distances) measures were calculated based on the rarefied OTU counts.

**Metagenomic sequencing**

Samples were metagenomically sequenced as one library each multiplexed through Illumina HiSeq machines and sequenced using the 2 × 100 bp paired-end read protocol. PRINSEQ v0.20.4 (*44*) was employed to sample dereplication and low complexity filtering. The length of each reads was trimmed with FASTX from the 5′ e and 3′ end using a quality threshold of 20. Read pairs with either reads was shorter than 60 bp or contained “N” were removed. 3) deduplicate the reads. Bowtie2 v2.2.5 (*45*) (using --reorder --no-contain --dovetail) was used to map reads to the human genome for decontamination.

**Taxonomy analysis**

Taxonomic profiling of the metagenomic samples was performed using MetaPhlAn2 v2.6.02, which uses a library of clade-specific markers to provide pan-microbial (bacterial, archaeal, viral and eukaryotic) quantification at the species level. MetaPhlAn2 (*46*) was run using default settings.

**Metabolomics profiling of human** **serum samples**

For the discovery cohort and external validation cohort1, targeted identification and quantification of serum metabolites was performed using an ultra-performance liquid chromatography coupled to tandem mass spectrometry (UPLC-MS/MS) system. This platform provides measures of 199 serum metabolome traits, including 12 subclasses (Table S14).

All of the standards of targeted metabolites were commercially purchased from Sigma-Aldrich (St. Louis, MO, USA), Steraloids Inc. (Newport, RI, USA) and TRC Chemicals (Toronto, ON, Canada). All the standards were prepared in water, methanol, sodium hydroxide solution, or hydrochloric acid solution to obtain individual stock solution at a concentration of 5.0 mg/mL. Appropriate amount of each stock solution was mixed to create stock calibration solutions.

Samples were thawed on ice-bath to diminish sample degradation and prepared as follows: 25μL of plasma was added to a 96-well plate and then the plate was transferred to the Biomek 4000 workstation (Biomek 4000, Beckman Coulter, Inc., Brea, California, USA). Three types of quality control samples i.e., test mixtures, internal standards, and pooled biological samples are routinely used in metabolomics platform. In addition to the quality controls, conditioning samples, and solvent blank samples are also required for obtaining optimal instrument performance. 100μL ice cold methanol with partial internal standards was automatically added to each sample and vortexed vigorously for 5 minutes. The plate was centrifuged at 4000g for 30 minutes (Allegra X-15R, Beckman Coulter, Inc., Indianapolis, IN, USA). Then the plate was returned back to the workstation. 30μL of supernatant was transferred to a clean 96-well plate, and 20μL of freshly prepared derivative reagents was added to each well. The plate was sealed and the derivatization was carried out at 30°C for 60 min. After derivatization, 350μL of ice-cold 50% methanol solution was added to dilute the sample. Then the plate was stored at -20°C for 20 minutes and followed by 4000g centrifugation at 4 °C for 30 minutes. 135μL of supernatant was transferred to a new 96-well plate with 15μL internal standards in each well. Serial dilutions of derivatized stock standards were added to the left wells. Finally the plate was sealed for LC-MS analysis. The raw data files from UPLC-MS/MS were processed using the QuanMET software (v2.0, Metabo-Profile, Shanghai, China) to perform peak integration, calibration, and quantitation for each metabolite.

**T2D risk variants and genetic risk score**

We used 28 significant variants identified in a meta-analysis of CKB and AGEN-T2D studies (*47*) to construct a T2D genetic risk score(GRS) as

$${GRS}_{i}=\sum_{j=1}^{m} x_{ij}b_{j}$$

Where,${GRS}_{i}$ is a genetic risk score for individual *i,* *m* is the number of SNPs in the score, $x_{ij}$ represented the number of the risk allele on two chromosomes for *ith* individual and *jth* SNP,$x_{ij}\in\{0,1,2\}$,$b_{j}$ represent the natural logarithm of the published odds ratio.

**Detailed machine learning parameters**

For the LightGBM method, the core parameters were set as follows: 'metric' : 'auc', 'boosting_type' : 'gbdt', 'colsample_bytree' : 0.9234, 'num_leaves' : 13, 'max_depth' : -1, 'n_estimators' : 200, 'min_child_samples': 399, 'min_child_weight': 0.1, 'reg_alpha': 2, 'reg_lambda': 5, 'subsample': 0.855. num_round: 10000, early_stopping_rounds: 50, verbose_eval: 1000.

**Fecal suspension inoculum preparation**

Each fecal sample (0.5 g) was diluted in 5 mL of a 0.09% (w/v) sterile normal saline in an anaerobic chamber (80% N_2_:10% CO_2_:10% H_2_). The fecal material was suspended by thorough vortexing (5 min) and centrifuged at 4 °C 300 rpm/min for 5 min. The clarified supernatant was transferred to a clean tube and used immediately for gut microbiota transplantation. Surveillance for bacterial contamination was performed by periodic bacteriological examinations of feces, food and padding. Normal saline was added into the samples with sufficient mixing. The mixtures were then cultured using the spread plate method on: 1) LB agar, Brain Heart Infusion agar and Thioglycolate agar under aerobic condition at 37°C for aerobic bacteria; 2) on Gifu anaerobic medium (GAM) agar under anaerobic condition at 37°C for anaerobic bacteria; and 3) on Modified Martin Agar and Tryptone Soya agar under aerobic condition at 25-28°C for fungi. All cultures were examined under optical microscope after 1, 2, 4, 7 and 14 days.

**Data availability.** For the discovery and external validation cohort 1, the raw data for 16 S rRNA gene sequences are available in the CNSA (https://db.cngb.org/cnsa/) of CNGBdb at accession number CNP0000829. For the external validation cohort 2, the raw data for 16 S rRNA gene sequences are available from the European Nucleotide Archive (https://www.ebi.ac.uk/ena/) at accession number PRJEB18535.

**Fig.S1. Overview of the discovery cohort: Guangzhou Nutrition and Health Study**

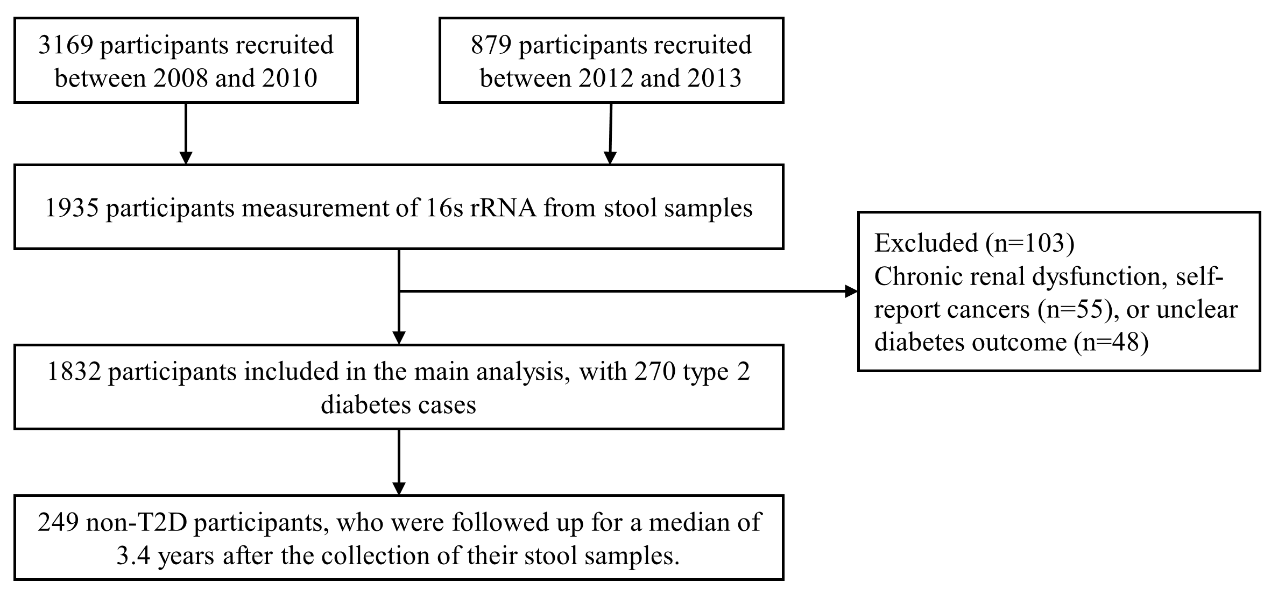

**Fig.S2. Compare the impact of the selected features on the model output in the discovery cohort.** We quantify the impacts of each feature on the model output using SHAP values, and plot the SHAP values of each feature for all samples. The color represents the feature value (red: higher level, blue: lower level), SHAP value greater than zero indicates that the feature may increase the type 2 diabetes risk for the given sample, otherwise, decrease the type 2 diabetes risk.

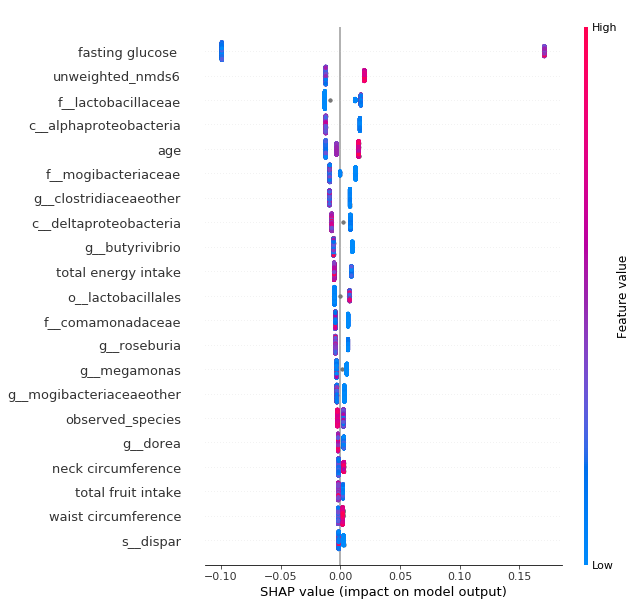

**Fig.S3.The marginal effect of individual selected features on type 2 diabetes.** We plot the SHAP values of every feature for each sample. X-axis represents the feature variable, while Y-axis represents the SHAP value for the feature variable. SHAP value greater than zero indicates that the feature may increase the type 2 diabetes risk for the given sample, otherwise, decrease the disease risk.

| 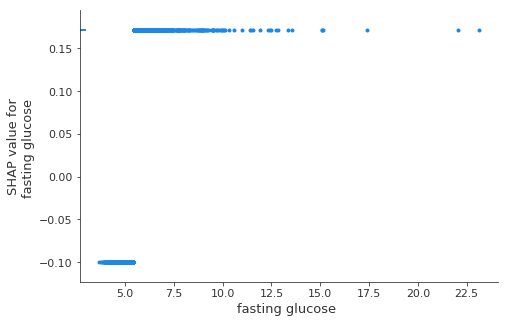 | 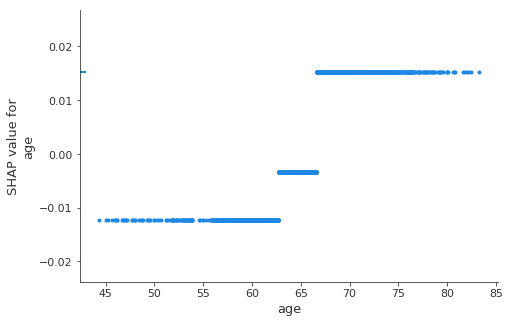 |
| --- | --- |
| 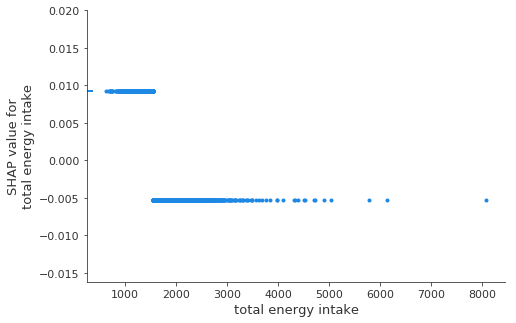 | 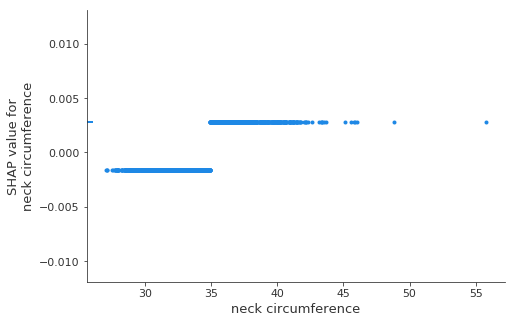 |
| 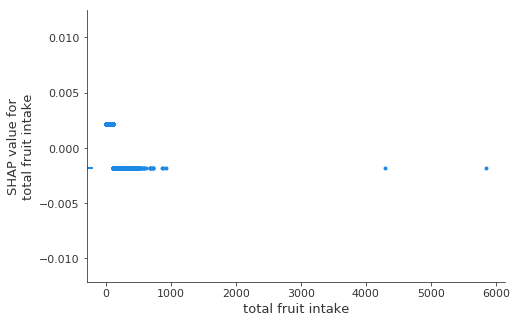 | 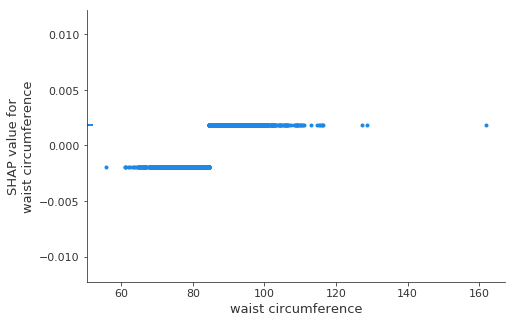 |

| 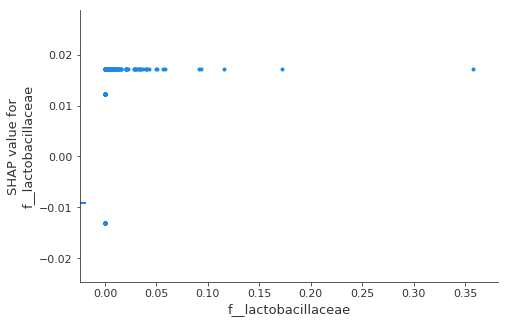 | 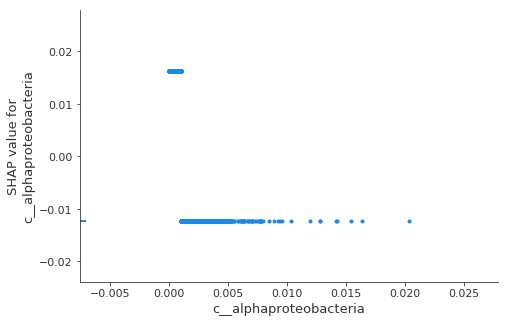 |
| --- | --- |
| 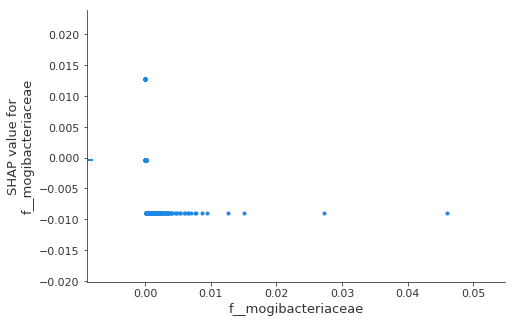 | 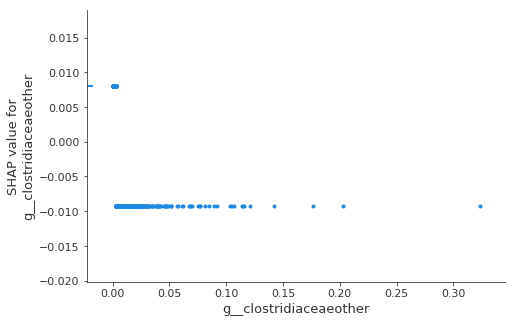 |
| 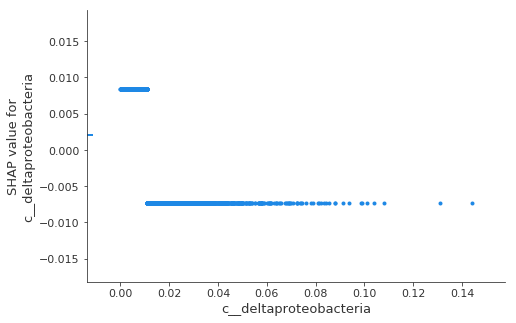 | 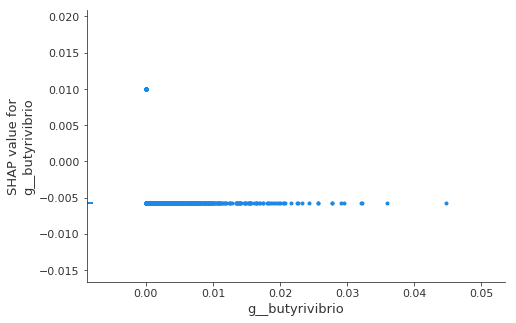 |
| 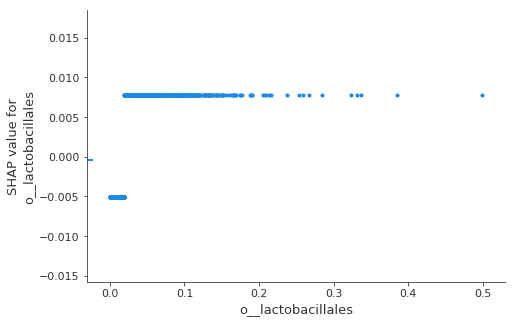 | 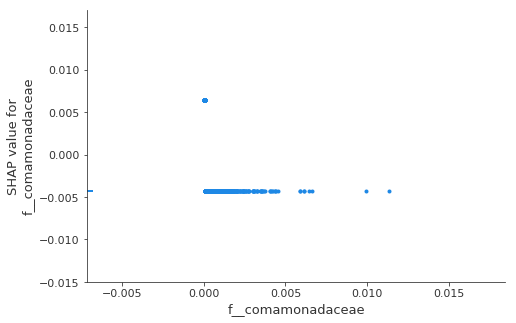 |
| 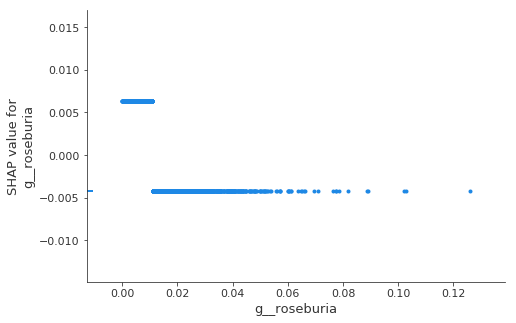 | 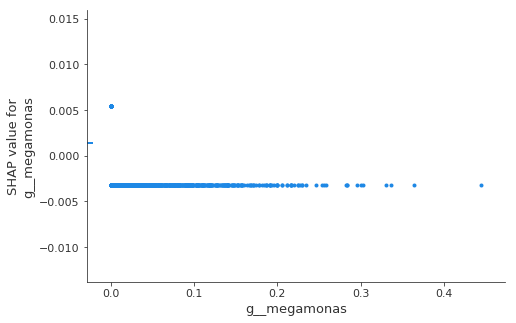 |
| 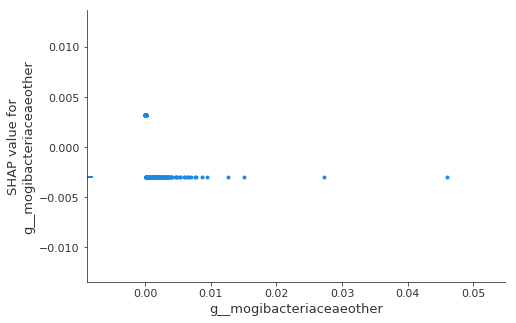 | 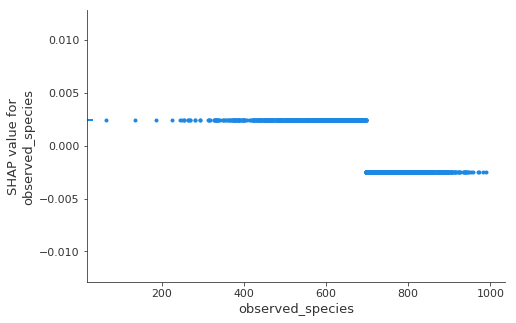 |
| 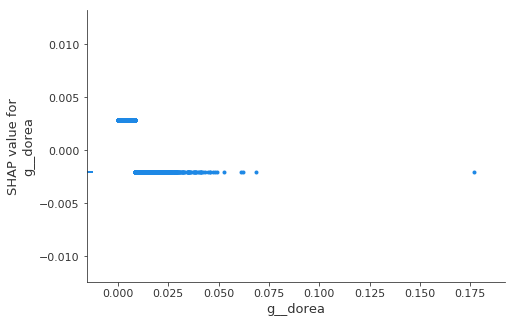 | 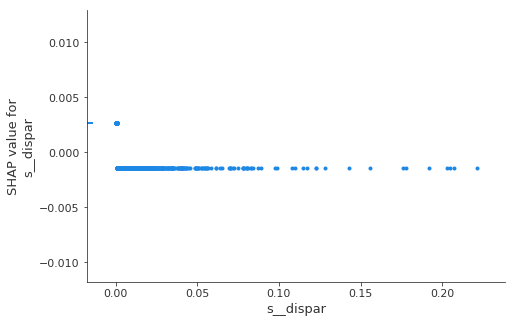 |

**Fig.S4. Associations of the selected microbiome features with risk of type 2 diabetes (T2D).** In this graph, we only present the microbiome that was significantly associated with T2D risk, other results are presented in Table S6. (A) Multivariable Poisson regression model was used to examine the association with T2D for each selected taxa-related features at higher abundance (i.e., higher the optimal threshold) with those at lower abundance (i.e., lower the optimal threshold). Covariates included in the statistical models for the discovery cohort and external validation cohort 1 were as follows: age, sex, BMI, waist circumference, total energy intake, alcohol drinking, smoking, household income, marital status, and self-reported educational level. For external validation cohort 2, all aforementioned covariates but total energy intake (not collected in external validation cohort 2) were used in the statistical model. (B) Multivariable Poisson regression model was used to estimate T2D risk per SD change in the selected taxa-related features, adjusted for the abovementioned covariates.

**A.**

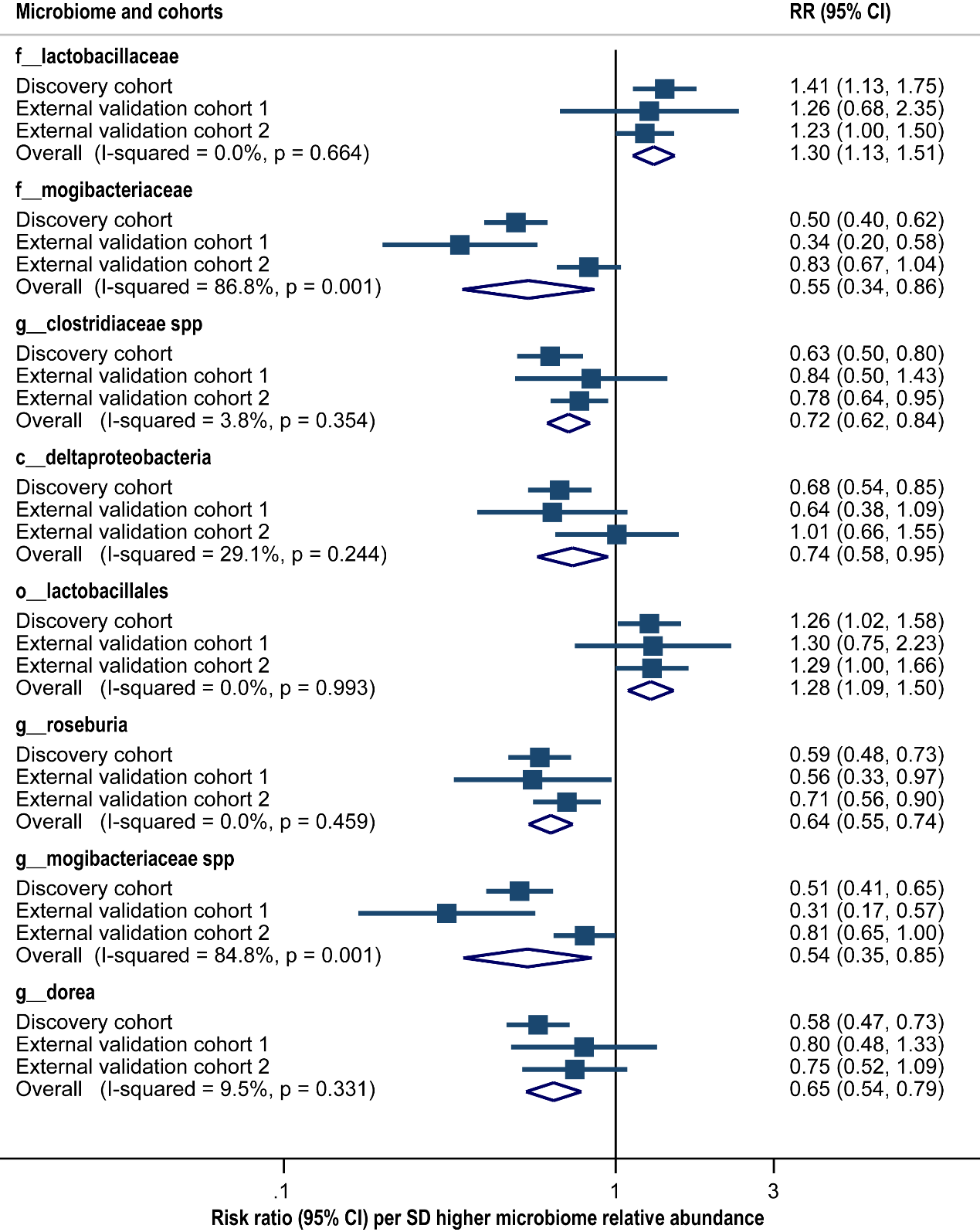

**B.**

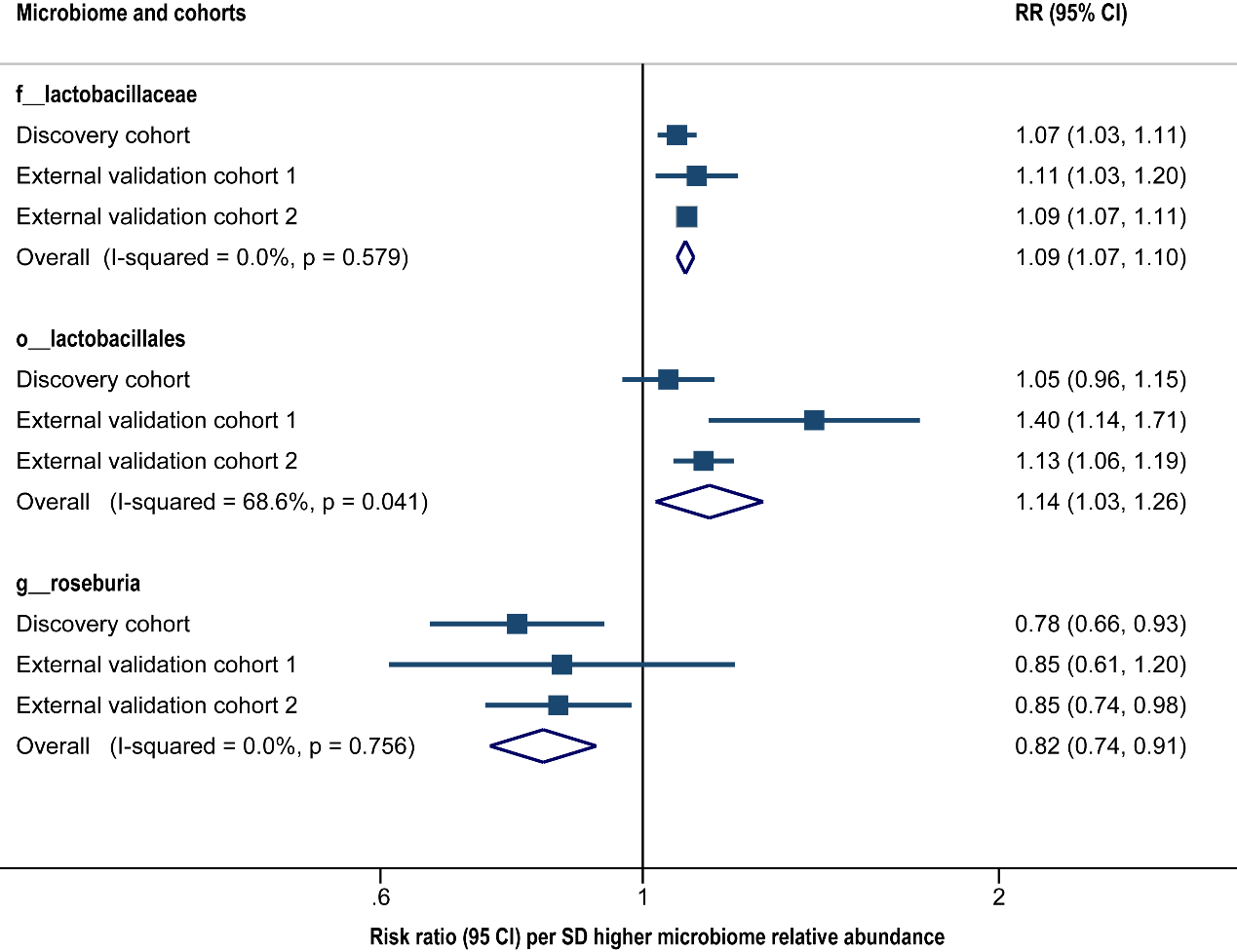

**Fig.S5. The mice weight of each group during follow-up. (A)** Body weight curves. **(B)** Quantification of body weight by AUC. Shown are mean± SD. * compared with CON group, # compared with Low MRS group, + compared with High MRS+non-T2D group. (*, #, +) *P*< 0.05, (**, ##, ++) *P*< 0.01, (***, ###, +++) *P*< 0.001 by ANOVA. The *P*-values were adjusted using the Benjamini and Hochberg method.

**A.**

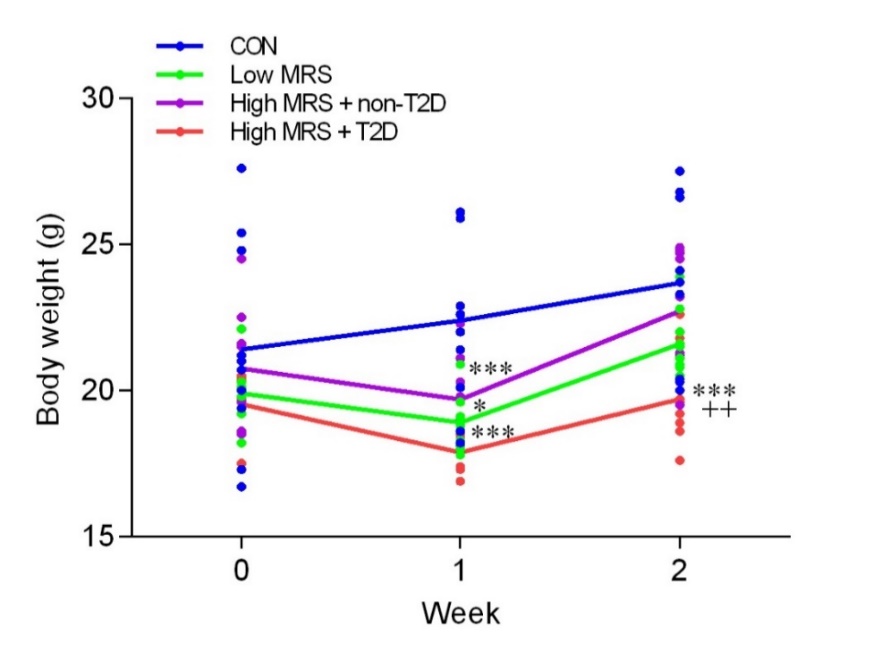

**B.**

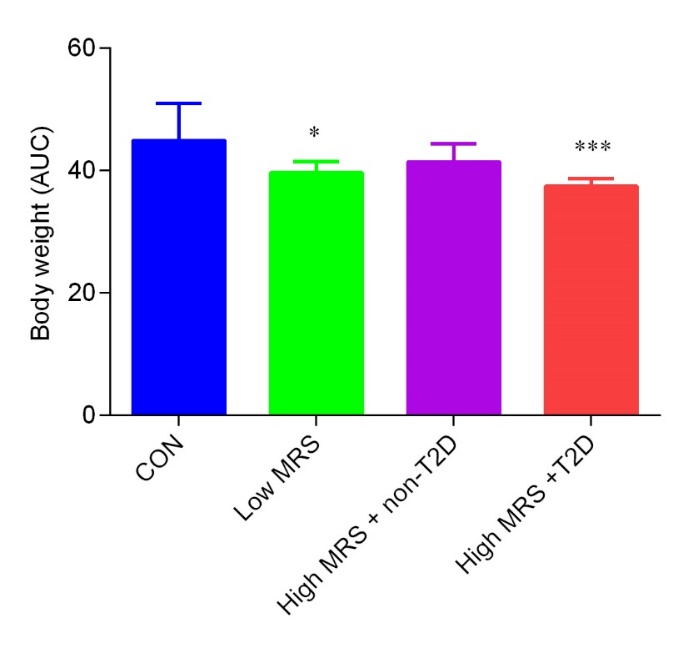

**Table S1. Comparison of the prediction performance of all inputed and selected features in different cohorts***

| **Features** | **Cohorts** | | | | | |
| --- | --- | --- | --- | --- | --- | --- |
|  | Internal validation cohort | | Internal test cohort | | External validation cohort 1 | |
|  | AUC | 95% CI | AUC | 95% CI | AUC | 95% CI |
| All | 0.89 | 0.84-0.92 | 0.87 | 0.83-0.92 | 0.86 | 0.80-0.91 |
| Selected | 0.88 | 0.84-0.92 | 0.87 | 0.82-0.92 | 0.83 | 0.77-0.88 |

*To train and validate our model, we divided the discovery cohort into three parts randomly with the ratio of 6:2:2 (training cohort, internal validation cohort, and internal test cohort). We further verified the capability of the model in an independent external validation cohort. Selected features (n=21) from the model showed a similar predictive capacity compared to all input features (n=297).

AUC: area under the ROC curve; CI: confidence interval.

**Table S2. Comparison of the prediction performance of LightGBM and random forest in different cohorts**

| **Algorithm** | **Internal validation** | | | **External validation** |
| --- | --- | --- | --- | --- |
|  | AUC (mean) | AUC (minimum) | AUC (maximum) | AUC |
| LightGBM | 0.91 | 0.85 | 0.98 | 0.72 |
| Random forest | 0.84 | 0.79 | 0.88 | 0.53 |

**Table S3. The mean absolute value of SHAP values for selected features***

| **Selected features** | **Mean absolute value of SHAP values** |
| --- | --- |
| fasting glucose | 0.126 |
| unweighted_nmds6 | 0.0153 |
| f__lactobacillaceae | 0.0144 |
| c__alphaproteobacteria | 0.0140 |
| age | 0.0109 |
| f__mogibacteriaceae | 0.00931 |
| g__clostridiaceae spp | 0.00854 |
| c__deltaproteobacteria | 0.0078 |
| g__butyrivibrio | 0.0073 |
| total energy intake | 0.00677 |
| o__lactobacillales | 0.00607 |
| f__comamonadaceae | 0.00519 |
| g__roseburia | 0.00503 |
| g__megamonas | 0.00406 |
| g__mogibacteriaceae spp | 0.00312 |
| observed_species | 0.00247 |
| g__dorea | 0.00237 |
| neck circumference | 0.00201 |
| fruit | 0.00197 |
| waist circumference | 0.00191 |
| s__dispar | 0.00188 |

*We took the mean absolute value of SHAP values for the selected features to get their average impact on T2D.

**Table S4. The inter-correlation of the selected microbiome features in the discovery cohort and external validation cohort 1**

| **Discovery cohort** |  | f__lactobacillaceae | c__alphaproteobacteria | f__mogibacteriaceae | g__clostridiaceae spp | c__deltaproteobacteria | g__butyrivibrio | o__lactobacillales | f__comamonadaceae | g__roseburia | g__megamonas | g__mogibacteriaceae spp | g__dorea | s__dispar |
| --- | --- | --- | --- | --- | --- | --- | --- | --- | --- | --- | --- | --- | --- | --- |
|  | f__lactobacillaceae | 1.000 | -0.067 | 0.070 | 0.037 | -0.041 | -0.023 | 0.370 | -0.042 | -0.065 | -0.046 | 0.070 | -0.076 | 0.075 |
|  | c__alphaproteobacteria | -0.067 | 1.000 | 0.020 | -0.056 | 0.009 | -0.017 | -0.041 | 0.046 | 0.180 | -0.011 | 0.020 | 0.341 | -0.080 |
|  | f__mogibacteriaceae | 0.070 | 0.020 | 1.000 | 0.388 | 0.168 | 0.000 | 0.057 | -0.053 | -0.014 | -0.006 | 1.000 | 0.008 | 0.133 |
|  | g__clostridiaceae spp | 0.037 | -0.056 | 0.388 | 1.000 | 0.218 | -0.012 | 0.310 | -0.093 | -0.043 | -0.038 | 0.388 | -0.017 | 0.159 |
|  | c__deltaproteobacteria | -0.041 | 0.009 | 0.168 | 0.218 | 1.000 | 0.091 | -0.023 | 0.101 | -0.086 | -0.144 | 0.168 | -0.033 | -0.118 |
|  | g__butyrivibrio | -0.023 | -0.017 | 0.000 | -0.012 | 0.091 | 1.000 | 0.013 | -0.087 | 0.075 | 0.009 | 0.000 | 0.371 | -0.034 |
|  | o__lactobacillales | 0.370 | -0.041 | 0.057 | 0.310 | -0.023 | 0.013 | 1.000 | -0.085 | -0.023 | -0.094 | 0.057 | -0.023 | 0.211 |
|  | f__comamonadaceae | -0.042 | 0.046 | -0.053 | -0.093 | 0.101 | -0.087 | -0.085 | 1.000 | -0.081 | -0.102 | -0.053 | -0.072 | -0.111 |
|  | g__roseburia | -0.065 | 0.180 | -0.014 | -0.043 | -0.086 | 0.075 | -0.023 | -0.081 | 1.000 | 0.003 | -0.014 | 0.529 | -0.029 |
|  | g__megamonas | -0.046 | -0.011 | -0.006 | -0.038 | -0.144 | 0.009 | -0.094 | -0.102 | 0.003 | 1.000 | -0.006 | 0.011 | -0.097 |
|  | g__mogibacteriaceae spp | 0.070 | 0.020 | 1.000 | 0.388 | 0.168 | 0.000 | 0.057 | -0.053 | -0.014 | -0.006 | 1.000 | 0.008 | 0.133 |
|  | g__dorea | -0.076 | 0.341 | 0.008 | -0.017 | -0.033 | 0.371 | -0.023 | -0.072 | 0.529 | 0.011 | 0.008 | 1.000 | -0.054 |
|  | s__dispar | 0.075 | -0.080 | 0.133 | 0.159 | -0.118 | -0.034 | 0.211 | -0.111 | -0.029 | -0.097 | 0.133 | -0.054 | 1.000 |
| **External validation cohort 1** |  | f__lactobacillaceae | c__alphaproteobacteria | f__mogibacteriaceae | g__clostridiaceae spp | c__deltaproteobacteria | g__butyrivibrio | o__lactobacillales | f__comamonadaceae | g__roseburia | g__megamonas | g__mogibacteriaceae spp | g__dorea | s__dispar |
|  | f__lactobacillaceae | 1.000 | -0.120 | -0.040 | 0.004 | -0.050 | 0.069 | 0.280 | -0.074 | -0.117 | -0.053 | -0.040 | -0.097 | -0.038 |
|  | c__alphaproteobacteria | -0.120 | 1.000 | -0.110 | -0.174 | 0.080 | -0.071 | -0.226 | 0.136 | 0.270 | -0.102 | -0.110 | 0.324 | -0.072 |
|  | f__mogibacteriaceae | -0.040 | -0.110 | 1.000 | 0.346 | 0.118 | -0.015 | 0.195 | -0.088 | 0.049 | -0.076 | 1.000 | -0.030 | 0.209 |
|  | g__clostridiaceae spp | 0.004 | -0.174 | 0.346 | 1.000 | 0.345 | -0.018 | 0.307 | -0.113 | -0.029 | -0.043 | 0.346 | -0.026 | 0.399 |
|  | c__deltaproteobacteria | -0.050 | 0.080 | 0.118 | 0.345 | 1.000 | -0.038 | -0.060 | 0.174 | -0.154 | -0.148 | 0.118 | -0.053 | 0.144 |
|  | g__butyrivibrio | 0.069 | -0.071 | -0.015 | -0.018 | -0.038 | 1.000 | -0.016 | -0.102 | 0.058 | -0.100 | -0.015 | 0.326 | -0.056 |
|  | o__lactobacillales | 0.280 | -0.226 | 0.195 | 0.307 | -0.060 | -0.016 | 1.000 | -0.111 | -0.136 | 0.048 | 0.195 | -0.154 | 0.078 |
|  | f__comamonadaceae | -0.074 | 0.136 | -0.088 | -0.113 | 0.174 | -0.102 | -0.111 | 1.000 | -0.103 | -0.097 | -0.088 | -0.057 | -0.067 |
|  | g__roseburia | -0.117 | 0.270 | 0.049 | -0.029 | -0.154 | 0.058 | -0.136 | -0.103 | 1.000 | -0.132 | 0.049 | 0.732 | 0.078 |
|  | g__megamonas | -0.053 | -0.102 | -0.076 | -0.043 | -0.148 | -0.100 | 0.048 | -0.097 | -0.132 | 1.000 | -0.076 | -0.163 | -0.056 |
|  | g__mogibacteriaceae spp | -0.040 | -0.110 | 1.000 | 0.346 | 0.118 | -0.015 | 0.195 | -0.088 | 0.049 | -0.076 | 1.000 | -0.030 | 0.209 |
|  | g__dorea | -0.097 | 0.324 | -0.030 | -0.026 | -0.053 | 0.326 | -0.154 | -0.057 | 0.732 | -0.163 | -0.030 | 1.000 | -0.032 |
|  | s__dispar | -0.038 | -0.072 | 0.209 | 0.399 | 0.144 | -0.056 | 0.078 | -0.067 | 0.078 | -0.056 | 0.209 | -0.032 | 1.000 |

**Table S5. The optimal threshold of the selected microbiome features according to their SHAP dependence plot**

| **Microbiome** | **Optimal threshold (relative abundance)** | **Taxa annotation** |
| --- | --- | --- |
| f__lactobacillaceae | 0.0000877 | p__Firmicutes; c__Bacilli; o__Lactobacillales; f__lactobacillaceae |
| c__alphaproteobacteria | 0.00101 | p__Proteobacteria; c__alphaproteobacteria |
| f__mogibacteriaceae | 0.0000403 | p__Firmicutes; c__Clostridia; o__Clostridiales; f__mogibacteriaceae |
| g__clostridiaceae spp | 0.00313 | p__Firmicutes; c__Clostridia; o__Clostridiales; f__Clostridiaceae; g__ |
| c__deltaproteobacteria | 0.0109 | p__Proteobacteria; c__deltaproteobacteria |
| g__butyrivibrio | 0.0000448 | p__Firmicutes; c__Clostridia; o__Clostridiales; f__Lachnospiraceae; g__butyrivibrio |
| o__lactobacillales | 0.0193 | p__Firmicutes; c__Bacilli; o__lactobacillales |
| f__comamonadaceae | 0.0000645 | p__Proteobacteria; c__Betaproteobacteria; o__Burkholderiales; f__comamonadaceae |
| g__roseburia | 0.011 | p__Firmicutes; c__Clostridia; o__Clostridiales; f__Lachnospiraceae; g__roseburia |
| g__megamonas | 0.00054 | p__Firmicutes; c__Clostridia; o__Clostridiales; f__Veillonellaceae; g__megamonas |
| g__mogibacteriaceae spp | 0.0000855 | p__Firmicutes; c__Clostridia; o__Clostridiales; f__mogibacteriaceae; g__ |
| g__dorea | 0.00861 | p__Firmicutes; c__Clostridia; o__Clostridiales; f__Lachnospiraceae; g__dorea |
| s__dispar | 0.000757 | p__Firmicutes; c__Clostridia; o__Clostridiales; f__Veillonellaceae; g__Veillonella; s__dispar |

**Table S6. Associations of the identified microbiome features with risk of type 2 diabetes (T2D) in different cohorts***

| **Microbiome was treated as binary variable** | | | | | | | **Microbiome was treated as continuous variable** | | | | | |
| --- | --- | --- | --- | --- | --- | --- | --- | --- | --- | --- | --- | --- |
| Identified taxa-related features | Cohorts | | | | | | Cohorts | | | | | |
|  | Discovery cohort | | External validation cohort 1 | | External validation cohort 2 | | Discovery cohort | | External validation cohort 1 | | External validation cohort 2 | |
|  | rr | 95% CI | rr | 95% CI | rr | 95% CI | rr | 95% CI | rr | 95% CI | rr | 95% CI |
| f__lactobacillaceae | 1.41 | 1.13-1.75 | 1.26 | 0.68-2.35 | 1.23 | 1-1.5 | 1.07 | 1.03-1.11 | 1.11 | 1.03-1.2 | 1.09 | 1.07-1.11 |
| c__alphaproteobacteria | 0.60 | 0.48-0.75 | 0.69 | 0.41-1.15 | 1.04 | 0.79-1.37 | 0.75 | 0.63-0.89 | 1.02 | 0.73-1.44 | 1.11 | 1.06-1.16 |
| f__mogibacteriaceae | 0.50 | 0.40-0.62 | 0.34 | 0.2-0.58 | 0.83 | 0.67-1.04 | 0.89 | 0.69-1.16 | 0.87 | 0.62-1.22 | 0.98 | 0.89-1.08 |
| g__clostridiaceae spp | 0.63 | 0.50-0.80 | 0.84 | 0.5-1.43 | 0.78 | 0.64-0.95 | 0.79 | 0.62-1.01 | 0.96 | 0.75-1.23 | 0.96 | 0.86-1.07 |
| c__deltaproteobacteria | 0.68 | 0.54-0.85 | 0.64 | 0.38-1.09 | 1.01 | 0.66-1.55 | 0.88 | 0.76-1.03 | 0.73 | 0.52-1.03 | 1.02 | 0.92-1.15 |
| g__butyrivibrio | 0.67 | 0.54-0.83 | 0.70 | 0.43-1.16 | NA | NA | 0.83 | 0.66-1.04 | 0.93 | 0.73-1.18 | NA | NA |
| o__lactobacillales | 1.26 | 1.02-1.58 | 1.30 | 0.75-2.23 | 1.29 | 1-1.66 | 1.05 | 0.96-1.15 | 1.40 | 1.14-1.71 | 1.13 | 1.06-1.19 |
| f__comamonadaceae | 0.61 | 0.48-0.76 | 0.75 | 0.44-1.27 | 1.28 | 0.92-1.78 | 0.94 | 0.78-1.12 | 0.74 | 0.52-1.07 | 0.99 | 0.92-1.07 |
| g__roseburia | 0.59 | 0.48-0.73 | 0.56 | 0.33-0.97 | 0.71 | 0.56-0.9 | 0.78 | 0.66-0.93 | 0.85 | 0.61-1.2 | 0.85 | 0.74-0.98 |
| g__megamonas | 0.66 | 0.53-0.82 | 0.72 | 0.44-1.18 | 1.04 | 0.85-1.28 | 1.05 | 0.94-1.16 | 0.70 | 0.43-1.12 | 1.04 | 0.95-1.13 |
| g__mogibacteriacea spp | 0.51 | 0.41-0.65 | 0.31 | 0.17-0.57 | 0.81 | 0.65-1 | 0.89 | 0.69-1.16 | 0.87 | 0.62-1.22 | 0.98 | 0.88-1.09 |
| g__dorea | 0.58 | 0.47-0.73 | 0.80 | 0.48-1.33 | 0.75 | 0.52-1.08 | 0.70 | 0.59-0.83 | 0.97 | 0.74-1.28 | 0.97 | 0.85-1.1 |
| s__dispar | 0.68 | 0.55-0.84 | 0.77 | 0.47-1.27 | 0.97 | 0.78-1.2 | 1.02 | 0.91-1.14 | 0.70 | 0.44-1.09 | 1.04 | 0.97-1.11 |

* Poisson regression model was used to examine the association with T2D for each identified taxa-related features. Covariates included in the statistical models for discovery cohort and external validation cohort 1 were as follows: age, sex, BMI, waist circumference, total energy intake, alcohol drinking, smoking, household income, marital status, and self-reported educational level. For external validation cohort 2, all aforementioned covariates but total energy intake (not collected in external validation cohort 2) were used in the statistical model.

rr: risk ratio; CI: confidence interval

**Table S7. Comparison of the model prediction performance in different cohorts based on selected microbiome features, host genetics, lifestyle and diet, traditional risk factors, and their combination***

| **Features** | **Cohorts** | | | | | |
| --- | --- | --- | --- | --- | --- | --- |
|  | Internal validation cohort | | Internal test cohort | | External validation cohort 1 | |
|  | AUC | 95% CI | AUC | 95% CI | AUC | 95% CI |
| Host genetics | 0.56 | 0.47-0.64 | 0.58 | 0.50-0.66 | 0.49 | 0.41-0.57 |
| FORS+Lifestyle+Diet | 0.63 | 0.55-0.71 | 0.66 | 0.57-0.76 | 0.51 | 0.45-0.57 |
| Selected Microbiome features | 0.72 | 0.64-0.81 | 0.71 | 0.62-0.80 | 0.6 | 0.52-0.69 |
| FORS+lifestyle+Diet+Microbiome features | 0.73 | 0.66-0.80 | 0.73 | 0.65-0.82 | 0.64 | 0.56-0.71 |

*AUC: area under the ROC curve; CI: confidence interval.

**Table S8. Associations of the microbiome risk score with type 2 diabetes (T2D)***

| **Outcome** | **Exposure** | **Cohorts** | | | | | | | | | | | |
| --- | --- | --- | --- | --- | --- | --- | --- | --- | --- | --- | --- | --- | --- |
|  |  | Discovery cohort | | | | External validation cohort 1 | | | | External validation cohort 2 (GGMP) | | | |
|  |  | n | rr | 95% CI | p | n | rr | 95% CI | p | n | rr | 95% CI | p |
| T2D | Microbiome risk score# | 1807 | 1.28 | 1.23-1.33 | 8.29E-33 | 188 | 1.23 | 1.13-1.34 | 1.49E-06 | 4221 (6787**) | 1.12 (1.08**) | 1.06-1.18 (1.03-1.13**) | 0.00012(0.00093**) |
|  | Microbiome risk score## | 1068 | 1.33 | 1.17-1.51 | 0.000011 |  |  |  |  |  |  |  |  |

*Poisson regression was performed to examine the association of the microbiome risk score with T2D risk.

The covariates for the discovery cohort and validation cohort 1 were total energy intake, age, waist circumference, sex, BMI, alcohol status, smoking status, education, marital status and income.

For the validation cohort 2 (GGMP), covariates including age, waist circumference, sex, BMI, alcohol status, smoking status, education, marital status and income

**For sensitivity analysis in validation cohort 2 (GGMP), excluding income as a covarite.

rr: risk ratio; CI: confidence interval.

\# The 16S-based microbiome risk score. Components including index of α-diversity (observe species) and 13 taxa-related features (f__lactobacillaceae, c__alphaproteobacteria, f__mogibacteriaceae, g__clostridiaceaeother c__deltaproteobacteria, g__butyrivibrio, o__lactobacillales, f__comamonadaceae, g__roseburia, g__megamonas, g__mogibacteriaceaeother, g__dorea, s__dispar).

\## The shotgun metagenome-based microbiome risk score. Components including index of α-diversity (observe species) and 13 taxa-related features (f__lactobacillaceae, c__alphaproteobacteria, f__mogibacteriaceae, g__clostridiaceaeother c__deltaproteobacteria, g__butyrivibrio, o__lactobacillales, f__comamonadaceae, g__roseburia, g__megamonas, g__mogibacteriaceaeother, g__dorea, s__dispar).

**Table S9. Associations of the microbiome risk score with prospective glucose increments***

| **Outcome** | **Model** | **n** | **Microbiome risk score** | | |
| --- | --- | --- | --- | --- | --- |
|  |  |  | beta | 95% CI | p |
| Glucose increments | 1 | 249 | 0.038 | 0.0081-0.068 | 0.013 |
| Glucose increments | 2 |  | 0.035 | 0.0049-0.064 | 0.022 |

*Linear regression was performed to examine the prospective association of baseline microbiome risk score with glucose increments, adjusted for :

Model 1: total energy intake + age + waist circumference + sex + BMI + alcohol status + smoking status + education + marital status + income

Model 2: Model 1+baseline glucose

Microbiome risk score: components including index of α-diversity (observe species) and 13 taxa-related features (f__lactobacillaceae, c__alphaproteobacteria, f__mogibacteriaceae, g__clostridiaceaeother c__deltaproteobacteria, g__butyrivibrio, o__lactobacillales, f__comamonadaceae, g__roseburia, g__megamonas, g__mogibacteriaceaeother, g__dorea, s__dispar).

**Table S10. Associations of baseline** **adiposity and dietary factors with microbiome risk score***

| **Exposures** | **n** | **microbiome risk score** | | |
| --- | --- | --- | --- | --- |
|  |  | beta | 95% CI | p |
| Age | 1812 | 0.023 | 0.0026, 0.043 | 0.027 |
| Energy intake | 1812 | 0.059 | -0.065, 0.18 | 0.35 |
| MET | 1812 | -0.02 | -0.12, 0.08 | 0.69 |
| BMI | 1812 | 0.1 | 0.023, 0.18 | 0.012 |
| Educatioin | 1812 | 0.2 | 0.042, 0.36 | 0.013 |
| Hip circumference | 1812 | -0.039 | -0.07, -0.007 | 0.017 |
| Waist circumference | 1812 | -0.0041 | -0.028, 0.02 | 0.74 |
| Neck circumference | 1812 | -0.037 | -0.099, 0.026 | 0.25 |
| Income | 1812 | -0.12 | -0.31, 0.06 | 0.19 |
| Red and processed meat intake | 1812 | -0.051 | -0.16, 0.59 | 0.37 |
| Fruit intake | 1812 | -0.025 | -0.14, 0.085 | 0.66 |
| Fish intake | 1812 | 0.061 | -0.046, 0.17 | 0.26 |
| Vegetable intake | 1812 | -0.08 | -0.19, 0.03 | 0.15 |
| Yogurt intake | 1812 | -0.027 | -0.13, 0.076 | 0.6 |
| Sex | 1812 | 0.035 | -0.38 0.45 | 0.87 |
| Current alcohol drinking | 1812 | -0.33 | -0.78, 0.12 | 0.15 |
| Current tea drinking | 1812 | -0.25 | -0.49, -0.018 | 0.035 |
| Current smoke drinking | 1812 | 0.09 | -0.3, 0.48 | 0.65 |
| Marital status | 1812 | 0.144 | -0.25, 0.54 | 0.47 |
| Drug use | 1812 | 2.56 | 2.18, 2.95 | <0.001 |

*beta: correlation coefficient of baseline diet and basic attributes with microbiome features; CI: confidence interval.

**Table S11. Associations of the microbiome risk score with body fat distribution in the discovery cohort***

| Outcome | | Microbiome risk score | | |
| --- | --- | --- | --- | --- |
|  |  | beta | 95% CI | p |
| TOTAL_FAT | 1750 | -5.344 | -27.28-16.59 | 0.63 |
| TOTAL_MASS | 1750 | -10.166 | -55.01-34.68 | 0.66 |
| TOTAL_PFAT | 1750 | -0.032 | -0.11-0.05 | 0.44 |
| ANDROID_FAT | 1750 | 2.577 | -7.22-12.38 | 0.61 |
| ANDROID_MASS | 1750 | 5.064 | -13.42-23.55 | 0.59 |
| ANDROID_PFAT | 1750 | 0.005 | -0.1-0.11 | 0.93 |
| GYNOID_FAT | 1750 | -7.921 | -21.4-5.56 | 0.25 |
| GYNOID_MASS | 1750 | -15.231 | -42.97-12.51 | 0.28 |
| GYNOID_PFAT | 1750 | -0.050 | -0.13-0.03 | 0.22 |
| TOTAL_PERCENT_FAT | 1750 | -0.004 | -0.08-0.08 | 0.92 |
| BODY_MASS_INDEX | 1750 | 0.149 | -0.45-0.74 | 0.62 |
| ANDROID_GYNOID_RATIO | 1750 | 0.002 | -0.00084-0.0047 | 0.17 |
| ANDROID_PERCENT_FAT | 1750 | 0.005 | -0.1-0.11 | 0.93 |
| GYNOID_PERCENT_FAT | 1750 | -0.050 | -0.13-0.03 | 0.22 |
| FAT_MASS_RATIO | 1750 | 0.005 | 0.0016-0.0074 | 0.00225 |
| TRUNK_LIMB_FAT_MASS_RATIO | 1750 | 0.007 | 0.0037-0.011 | 0.000117 |
| FAT_MASS_HEIGHT_SQUARED | 1750 | 0.033 | -0.05-0.11 | 0.422 |
| TOTAL_FAT_MASS | 1750 | 2.746 | -84.04-89.53 | 0.951 |
| GLOBAL_FAT | 1750 | -3.066 | -90.56-84.43 | 0.945 |
| GLOBAL_MASS | 1750 | -34.092 | -202.98-134.8 | 0.692 |
| GLOBAL_PFAT | 1750 | -0.016 | -0.1-0.07 | 0.705 |
| HEAD_FAT | 1750 | -0.368 | -2.12-1.38 | 0.681 |
| HEAD_MASS | 1750 | -2.770 | -10.15-4.62 | 0.462 |
| HEAD_PFAT | 1750 | 0.006 | -0.0026-0.0014 | 0.183 |
| LARM_FAT | 1750 | 1.654 | -4.96-8.27 | 0.624 |
| LARM_MASS | 1750 | -2.245 | -12.41-7.92 | 0.665 |
| LARM_PFAT | 1750 | 0.032 | -0.09-0.16 | 0.606 |
| RARM_FAT | 1750 | 2.092 | -4.3-8.48 | 0.521 |
| RARM_MASS | 1750 | -1.769 | -12.08-8.55 | 0.737 |
| RARM_PFAT | 1750 | 0.042 | -0.08-0.16 | 0.490 |
| TRUNK_FAT | 1750 | 26.380 | -24.08-76.84 | 0.306 |
| TRUNK_MASS | 1750 | 37.376 | -55.31-130.06 | 0.429 |
| TRUNK_PFAT | 1750 | 0.030 | -0.06-0.12 | 0.536 |
| L_LEG_FAT | 1750 | -14.150 | -29.21-0.91 | 0.066 |
| L_LEG_MASS | 1750 | -28.815 | -56.72--0.91 | 0.043 |
| L_LEG_PFAT | 1750 | -0.079 | -0.18-0.02 | 0.105 |
| R_LEG_FAT | 1750 | -14.513 | -30-0.97 | 0.066 |
| R_LEG_MASS | 1750 | -26.408 | -54.8-1.98 | 0.068 |
| R_LEG_PFAT | 1750 | -0.093 | -0.19-0.01 | 0.063 |
| SUBTOT_FAT | 1750 | 1.463 | -85.46-88.39 | 0.974 |
| SUBTOT_MASS | 1750 | -21.861 | -182.34-138.62 | 0.789 |
| SUBTOT_PFAT | 1750 | -0.007 | -0.09-0.08 | 0.868 |
| WBTOT_FAT | 1750 | 1.095 | -86.71-88.9 | 0.980 |
| WBTOT_MASS | 1750 | -24.630 | -189.48-140.21 | 0.770 |
| WBTOT_PFAT | 1750 | -0.006 | -0.09-0.08 | 0.888 |

*Linear regression was performed to examine the association of microbiome risk score with components of body fat distribution, adjusted for total energy intake, age, sex, alcohol status, smoking status, education, marital status and income

Microbiome risk score: components including index of α-diversity (observe species), and 13 taxa-related features (f__lactobacillaceae, c__alphaproteobacteria, f__mogibacteriaceae, g__clostridiaceaeother c__deltaproteobacteria, g__butyrivibrio, o__lactobacillales, f__comamonadaceae, g__roseburia, g__megamonas, g__mogibacteriaceaeother, g__dorea, s__dispar).

**Table S12. Associations of the microbiome risk score with body fat distribution in the external validation cohort 1***

| Outcome | n | Microbiome risk score | | |
| --- | --- | --- | --- | --- |
|  |  | Beta | 95% CI | p |
| TOTAL_FAT | 185 | -5.120 | -75.29-55.53 | 0.884 |
| TOTAL_MASS | 185 | 19.324 | -123.67-136.55 | 0.782 |
| TOTAL_PFAT | 185 | -0.102 | -0.35-0.15 | 0.449 |
| ANDROID_FAT | 185 | 15.973 | -12.86-41.36 | 0.273 |
| ANDROID_MASS | 185 | 37.384 | -18.1-83.13 | 0.169 |
| ANDROID_PFAT | 185 | 0.074 | -0.24-0.42 | 0.678 |
| GYNOID_FAT | 185 | -21.093 | -65.86-17.6 | 0.348 |
| GYNOID_MASS | 185 | -18.060 | -109.09-56.94 | 0.686 |
| GYNOID_PFAT | 185 | -0.191 | -0.44-0.06 | 0.157 |
| TOTAL_PERCENT_FAT | 185 | 0.009 | -0.23-0.26 | 0.943 |
| BODY_MASS_INDEX | 185 | 0.122 | -0.08-0.3 | 0.231 |
| ANDROID_GYNOID_RATIO | 185 | 0.009 | -0.00033-0.0178 | 0.059 |
| ANDROID_PERCENT_FAT | 185 | 0.074 | -0.24-0.42 | 0.678 |
| GYNOID_PERCENT_FAT | 185 | -0.191 | -0.44-0.06 | 0.157 |
| FAT_MASS_RATIO | 185 | 0.007 | 0.0067-0.016 | 0.159 |
| TRUNK_LIMB_FAT_MASS_RATIO | 185 | 0.015 | 0.0023-0.03 | 0.020 |
| FAT_MASS_HEIGHT_SQUARED | 185 | 0.045 | -0.06-0.15 | 0.438 |
| TOTAL_FAT_MASS | 185 | 102.950 | -177.6-360.91 | 0.477 |
| GLOBAL_FAT | 185 | 102.918 | -177.61-360.84 | 0.477 |
| GLOBAL_MASS | 185 | 213.248 | -313.55-684.25 | 0.427 |
| GLOBAL_PFAT | 185 | 0.009 | -0.23-0.26 | 0.944 |
| HEAD_FAT | 185 | 2.185 | -3.92-6.54 | 0.437 |
| HEAD_MASS | 185 | 4.644 | -20.15-23.75 | 0.694 |
| HEAD_PFAT | 185 | 0.021 | -0.01-0.04 | 0.108 |
| LARM_FAT | 185 | 4.281 | -15.06-23.13 | 0.677 |
| LARM_MASS | 185 | 9.323 | -20.45-36.65 | 0.544 |
| LARM_PFAT | 185 | -0.038 | -0.39-0.34 | 0.844 |
| RARM_FAT | 185 | 6.775 | -13.95-25.61 | 0.524 |
| RARM_MASS | 185 | 14.976 | -19.18-43.11 | 0.371 |
| RARM_PFAT | 185 | -0.028 | -0.37-0.35 | 0.885 |
| TRUNK_FAT | 185 | 103.849 | -39.88-242.11 | 0.171 |
| TRUNK_MASS | 185 | 202.665 | -76.77-457.47 | 0.158 |
| TRUNK_PFAT | 185 | 0.090 | -0.16-0.37 | 0.529 |
| L_LEG_FAT | 185 | -4.545 | -61.75-44.84 | 0.874 |
| L_LEG_MASS | 185 | -10.916 | -107.84-78.19 | 0.827 |
| L_LEG_PFAT | 185 | -0.075 | -0.42-0.23 | 0.669 |
| R_LEG_FAT | 185 | -9.819 | -65.59-40.44 | 0.731 |
| R_LEG_MASS | 185 | -8.410 | -102.94-75.64 | 0.861 |
| R_LEG_PFAT | 185 | -0.141 | -0.47-0.18 | 0.419 |
| SUBTOT_FAT | 185 | 100.541 | -176.34-356.24 | 0.483 |
| SUBTOT_MASS | 185 | 207.638 | -303.48-667.37 | 0.426 |
| SUBTOT_PFAT | 185 | 0.007 | -0.25-0.28 | 0.963 |
| WBTOT_FAT | 185 | 102.726 | -178.09-360.61 | 0.478 |
| WBTOT_MASS | 185 | 212.282 | -315.69-683.18 | 0.429 |
| WBTOT_PFAT | 185 | 0.010 | -0.23-0.26 | 0.942 |

*Linear regression was performed to examine the association of microbiome risk score with components of body fat distribution, adjusted for total energy intake, age, sex, alcohol status, smoking status, education, marital status and income

Microbiome risk score: components including index of α-diversity (observe species), and 13 taxa-related features (f__lactobacillaceae, c__alphaproteobacteria, f__mogibacteriaceae, g__clostridiaceaeother c__deltaproteobacteria, g__butyrivibrio, o__lactobacillales, f__comamonadaceae, g__roseburia, g__megamonas, g__mogibacteriaceaeother, g__dorea, s__dispar).

**Table S13. Risk ratio (95% CIs) of type 2 diabetes (T2D) according to tertiles of the microbiome risk score and trunk/limb fat mass ratio***

| Outcome | Exposure (T_MRS#T_Trunk/limb fat mass ratio ) | Discovery cohort | | | | External validation cohort1 | | | |
| --- | --- | --- | --- | --- | --- | --- | --- | --- | --- |
|  |  | n | rr | 95% CI | p | n | rr | 95% CI | p |
| T2D | 1 1(refrence) | 295 | 1 |  |  | 25 | 1 |  |  |
|  | 1 2 | 295 | 1.83 | 0.86-3.88 | 0.12 | 23 | 2.07 | 0.24-18.11 | 0.51 |
|  | 1 3 | 239 | 3.61 | 1.81-7.18 | 0.00026 | 17 | 6.12 | 0.95-39.64 | 0.057 |
|  | 2 1 | 156 | 2.04 | 0.89-4.69 | 0.093 | 27 | 4.59 | 0.7-29.85 | 0.11 |
|  | 2 2 | 144 | 3.02 | 1.40-6.54 | 0.005 | 29 | 4.76 | 0.77-29.52 | 0.094 |
|  | 2 3 | 164 | 5.7 | 2.91-11.16 | 3.91E-07 | 26 | 10.15 | 1.81-56.93 | 0.008 |
|  | 3 1 | 139 | 4.5 | 2.21-9.17 | 0.000035 | 10 | 10.16 | 1.49-69.35 | 0.018 |
|  | 3 2 | 151 | 6.14 | 3.12-12.08 | 1.48E-07 | 10 | 10.57 | 1.7-65.52 | 0.011 |
|  | 3 3 | 186 | 11.79 | 6.28-22.16 | 1.78E-14 | 18 | 15.32 | 2.7-86.88 | 0.002 |

*Poisson regression was performed to examine the interaction of the microbiome risk score with the trunk/limb fat mass ratio on T2D risk, adjusted for total energy intake, age, sex, alcohol status, smoking status, education, marital status and income

rr: risk ration; CI: confidence interval.

T_MRS: Tertile of microbiome risk score; T_Trunk/limb fat mass ratio: Tertile of Trunk/limb fat mass ratio.

**Table S14. Serum metabolite list measured by the ultra-performance liquid chromatography coupled to tandem mass spectrometry (UPLC-MS/MS) system**

| **Name** | **HMDB ID** | **Subclasses** |
| --- | --- | --- |
| 2_Hydroxy_2_methylbutyric acid | HMDB0001987 | Amino acids |
| 2-Methylbutyroylcarnitine | HMDB0000378 | Amino acids |
| 5_Aminolevulinic acid | HMDB0001149 | Amino acids |
| Acetylglycine | HMDB0000532 | Amino acids |
| Alpha_N_Phenylacetyl_L_glutamine | HMDB0006344 | Amino acids |
| Aminoadipic acid | HMDB0000510 | Amino acids |
| Beta_Alanine | HMDB0000056 | Amino acids |
| Citrulline | HMDB0000904 | Amino acids |
| Creatine | HMDB0000064 | Amino acids |
| Dimethylglycine | HMDB0000092 | Amino acids |
| Gamma_Aminobutyric acid | HMDB0000112 | Amino acids |
| Glycine | HMDB0000123 | Amino acids |
| Glycylproline | HMDB0000721 | Amino acids |
| Guanidoacetic acid | HMDB0000128 | Amino acids |
| L_Alanine | HMDB0000161 | Amino acids |
| L_Alpha_aminobutyric acid | HMDB0003911 | Amino acids |
| L_Arginine | HMDB0000517 | Amino acids |
| L_Asparagine | HMDB0000168 | Amino acids |
| L_Aspartic acid | HMDB0000191 | Amino acids |
| L_Cystine | HMDB0000192 | Amino acids |
| L_Glutamic acid | HMDB0000148 | Amino acids |
| L_Glutamine | HMDB0000641 | Amino acids |
| L_Histidine | HMDB0000177 | Amino acids |
| L_Homocitrulline | HMDB0000679 | Amino acids |
| L_Homoserine | HMDB0000719 | Amino acids |
| L_Leucine | HMDB0000687 | Amino acids |
| L_Lysine | HMDB0000182 | Amino acids |
| L_Methionine | HMDB0000696 | Amino acids |
| L_Phenylalanine | HMDB0000159 | Amino acids |
| L_Pipecolic acid | HMDB0000716 | Amino acids |
| L_Proline | HMDB0000162 | Amino acids |
| L_Serine | HMDB0000187 | Amino acids |
| L_threonine | HMDB0000167 | Amino acids |
| L_Tryptophan | HMDB0000929 | Amino acids |
| L_Tyrosine | HMDB0000158 | Amino acids |
| L_Valine | HMDB0000883 | Amino acids |
| Methionine sulfoxide | HMDB0002005 | Amino acids |
| Methylcysteine | HMDB0002108 | Amino acids |
| N_Acetyl_L_aspartic acid | HMDB0000812 | Amino acids |
| N_Acetylglutamine | HMDB0006029 | Amino acids |
| N_Acetylserine | HMDB0002931 | Amino acids |
| Norvaline | HMDB0013716 | Amino acids |
| Ornithine | HMDB0000214 | Amino acids |
| Pyroglutamic acid | HMDB0000267 | Amino acids |
| Sarcosine | HMDB0000271 | Amino acids |
| Selenomethionine | HMDB0003966 | Amino acids |
| 4_Hydroxyphenylpyruvic acid | HMDB0000707 | Benzenoids |
| Benzamide | HMDB0004461 | Benzenoids |
| Benzenebutanoic acid | HMDB0000543 | Benzenoids |
| Benzoic acid | HMDB0001870 | Benzenoids |
| Hippuric acid | HMDB0000714 | Benzenoids |
| Homovanillic acid | HMDB0000118 | Benzenoids |
| Mandelic acid | HMDB0000703 | Benzenoids |
| Ortho_Hydroxyphenylacetic acid | HMDB0000669 | Benzenoids |
| p_Hydroxyphenylacetic acid | HMDB0000020 | Benzenoids |
| Phenylacetic acid | HMDB0000209 | Benzenoids |
| Phenyllactic acid | HMDB0000779 | Benzenoids |
| Phenylpyruvic acid | HMDB0000205 | Benzenoids |
| Phthalic acid | HMDB0002107 | Benzenoids |
| Protocatechuic acid | HMDB0001856 | Benzenoids |
| Chenodeoxycholic acid | HMDB0000518 | Bile acids |
| Cholic acid | HMDB0000619 | Bile acids |
| Deoxycholic acid | HMDB0000626 | Bile acids |
| GLCA_3S | HMDB0002639 | Bile acids |
| Glycochenodeoxycholic acid | HMDB0000637 | Bile acids |
| Glycocholic acid | HMDB0000138 | Bile acids |
| Glycodeoxycholic acid | HMDB0000631 | Bile acids |
| Glycohyocholic acid |  | Bile acids |
| Glycoursodeoxycholic acid | HMDB0000708 | Bile acids |
| Hyodeoxycholic acid | HMDB0000733 | Bile acids |
| Taurochenodeoxycholic acid | HMDB0000951 | Bile acids |
| Taurocholic acid | HMDB0000036 | Bile acids |
| Taurohyodeoxycholic acid |  | Bile acids |
| Tauroursodeoxycholic acid | HMDB0000874 | Bile acids |
| D_Fructose | HMDB0000660 | Carbohydrates |
| D_Gluconolactone | HMDB0000150 | Carbohydrates |
| D_Glucose | HMDB0000122 | Carbohydrates |
| D_Maltose and Alpha_Lactose | HMDB0000163/HMDB0000186 | Carbohydrates |
| D_Xylose | HMDB0000098 | Carbohydrates |
| D_Xylulose | HMDB0001644 | Carbohydrates |
| Erythronic acid | HMDB0000613 | Carbohydrates |
| Fructose 1,6_bisphosphate | HMDB0001058 | Carbohydrates |
| Glucaric acid | HMDB0000663 | Carbohydrates |
| Glyceric acid | HMDB0000139 | Carbohydrates |
| N_Acetylneuraminic acid | HMDB0000230 | Carbohydrates |
| Phosphoribosyl pyrophosphate | HMDB0000280 | Carbohydrates |
| Rhamnose | HMDB0000849 | Carbohydrates |
| Tartaric acid | HMDB0000956 | Carbohydrates |
| Trehalose | HMDB0000975 | Carbohydrates |
| 3-Hydroxylisovalerylcarnitine | HMDB0061189 | Carnitines |
| Adipoylcarnitine | HMDB0061677 | Carnitines |
| Carnitine | HMDB0000062 | Carnitines |
| Decanoylcarnitine C10 | HMDB0000651 | Carnitines |
| Glutarylcarnitine | HMDB0013130 | Carnitines |
| Hexanylcarnitine | HMDB0000705 | Carnitines |
| Isovelarylcarnitine | HMDB0000688 | Carnitines |
| L-Acetylcarnitine | HMDB0000201 | Carnitines |
| Lauroylcarnitine | HMDB0002250 | Carnitines |
| Linoleylcarnitine | HMDB0006469 | Carnitines |
| Malonylcarnitine | HMDB0002095 | Carnitines |
| Methylmalonylcarnitine C4DC | HMDB0013133 | Carnitines |
| Myristoylcarnitine | HMDB0005066 | Carnitines |
| Octanoylcarnitine | HMDB0000791 | Carnitines |
| Oleylcarnitine C18 1 | HMDB0005065 | Carnitines |
| Palmitoylcarnitine | HMDB0000222 | Carnitines |
| Propionylcarnitine | HMDB0000824 | Carnitines |
| Stearylcarnitine | HMDB0000848 | Carnitines |
| 10Z_Heptadecenoic acid | HMDB0060038 | Fatty acids |
| 12_hydroxystearic acid | HMDB0061706 | Fatty acids |
| 2_Hydroxy_3_methylbutyric acid | HMDB0000407 | Fatty acids |
| 2_Hydroxycaproic acid | HMDB0001624 | Fatty acids |
| 3_Hydroxyisovaleric acid | HMDB0000754 | Fatty acids |
| 3_Methyladipic acid | HMDB0000555 | Fatty acids |
| 5_Dodecenoic acid | HMDB0000529 | Fatty acids |
| 8,11,14_Eicosatrienoic acid | HMDB0002925 | Fatty acids |
| Acetic acid | HMDB0000042 | Fatty acids |
| Adipic acid | HMDB0000448 | Fatty acids |
| Adrenic acid | HMDB0002226 | Fatty acids |
| Alpha_Linolenic acid | HMDB0001388 | Fatty acids |
| Arachidonic acid | HMDB0001043 | Fatty acids |
| Azelaic acid | HMDB0000784 | Fatty acids |
| Butyric acid | HMDB0000039 | Fatty acids |
| Caproic acid | HMDB0000535 | Fatty acids |
| Citramalic acid | HMDB0000426 | Fatty acids |
| Decanoic acid | HMDB0000511 | Fatty acids |
| Docosahexaenoic acid DHA | HMDB0002183 | Fatty acids |
| Docosapentaenoic acid 22n_6 | HMDB0001976 | Fatty acids |
| Docosapentaenoic acid DPA | HMDB0006528 | Fatty acids |
| Dodecanoic acid | HMDB0000638 | Fatty acids |
| Eicosapentaenoic acid EPA | HMDB0001999 | Fatty acids |
| Gamma_Linolenic acid | HMDB0003073 | Fatty acids |
| Heptadecanoic acid | HMDB0002259 | Fatty acids |
| Isocaproic acid | HMDB0000689 | Fatty acids |
| Isovaleric acid | HMDB0000718 | Fatty acids |
| Linoelaidic acid | HMDB0006270 | Fatty acids |
| Linoleic acid | HMDB0000673 | Fatty acids |
| Methylglutaric acid | HMDB0000752 | Fatty acids |
| Methylsuccinic acid | HMDB0001844 | Fatty acids |
| Myristelaidic acid | HMDB0062248 | Fatty acids |
| Myristic acid | HMDB0000806 | Fatty acids |
| Myristoleic acid | HMDB0002000 | Fatty acids |
| Nonadeca_10Z_enoic acid | HMDB0013622 | Fatty acids |
| Nonanoic acid | HMDB0000847 | Fatty acids |
| Octanoic acid | HMDB0000482 | Fatty acids |
| Oleic acid | HMDB0000207 | Fatty acids |
| Palmitelaidic acid | HMDB0012328 | Fatty acids |
| Palmitic acid | HMDB0000220 | Fatty acids |
| Palmitoleic acid | HMDB0003229 | Fatty acids |
| Pentadecanoic acid | HMDB0000826 | Fatty acids |
| Pimelic acid | HMDB0000857 | Fatty acids |
| Propanoic acid | HMDB0000237 | Fatty acids |
| Ricinoleic acid | HMDB0034297 | Fatty acids |
| Sebacic acid | HMDB0000792 | Fatty acids |
| Stearic acid | HMDB0000827 | Fatty acids |
| Suberic acid | HMDB0000893 | Fatty acids |
| Tridecanoic acid | HMDB0000910 | Fatty acids |
| Undecanoic acid | HMDB0000947 | Fatty acids |
| Undecylenic acid | HMDB0033724 | Fatty acids |
| 3_Indolepropionic acid | HMDB0002302 | Indoles |
| 5_Hydroxy_L_tryptophan | HMDB0000472 | Indoles |
| Indoleacetic acid | HMDB0000197 | Indoles |
| Adenosine monophosphate | HMDB0000045 | Nucleosides |
| 2_Hydroxybutyric acid | HMDB0000008 | Organic acids |
| 3_Hydroxybutyric acid | HMDB0000357 | Organic acids |
| 3_Methyl_2_oxovaleric acid | HMDB0000491 | Organic acids |
| Acetoacetic acid | HMDB0000060 | Organic acids |
| alpha_Hydroxyisobutyric acid | HMDB0000729 | Organic acids |
| Alpha_ketoisovaleric acid | HMDB0000019 | Organic acids |
| Cis and trans_Cinnamic acid |  | Organic acids |
| cis_Aconitic acid | HMDB0000072 | Organic acids |
| Citric acid | HMDB0000094 | Organic acids |
| D_2_Hydroxyglutaric acid | HMDB0000606 | Organic acids |
| Fumaric acid | HMDB0000134 | Organic acids |
| Galactonic acid | HMDB0000565 | Organic acids |
| Glutaconic acid | HMDB0000620 | Organic acids |
| Glutaric acid | HMDB0000661 | Organic acids |
| Glycolic acid | HMDB0000115 | Organic acids |
| Hydroxypropionic acid | HMDB0000700 | Organic acids |
| Isocitric acid | HMDB0000193 | Organic acids |
| Ketoleucine | HMDB0000695 | Organic acids |
| L_Lactic acid | HMDB0000190 | Organic acids |
| L_Malic acid | HMDB0000156 | Organic acids |
| Maleic acid | HMDB0000176 | Organic acids |
| Malonic acid | HMDB0000691 | Organic acids |
| Methylmalonic acid | HMDB0000202 | Organic acids |
| Oxalic acid | HMDB0002329 | Organic acids |
| Oxoadipic acid | HMDB0000225 | Organic acids |
| Oxoglutaric acid | HMDB0000208 | Organic acids |
| Pyruvic acid | HMDB0000243 | Organic acids |
| Succinic acid | HMDB0000254 | Organic acids |
| Threonic acid | HMDB0000943 | Organic acids |
| trans_Aconitic acid | HMDB0000958 | Organic acids |
| Shikimic acid | HMDB0003070 | Organooxygen compounds |
| 2_Phenylpropionate | HMDB0011743 | Phenylpropanoic acids |
| 3_ 3_Hydroxyphenyl _3_hydroxypropanoic acid | HMDB0002643 | Phenylpropanoic acids |
| Hydrocinnamic acid | HMDB0000764 | Phenylpropanoic acids |
| Hydroxyphenyllactic acid | HMDB0000755 | Phenylpropanoic acids |
| N_Methylnicotinamide | HMDB0003152 | Pyridines |
